# Proximity-dependent recruitment of Polycomb Repressive Complexes by the lncRNA *Airn*

**DOI:** 10.1101/2022.12.20.521198

**Authors:** Aki K. Braceros, Megan D. Schertzer, Arina Omer, Jackson B. Trotman, Eric S. Davis, Jill M. Dowen, Douglas H. Phanstiel, Erez Lieberman Aiden, J. Mauro Calabrese

## Abstract

During mouse embryogenesis, expression of the lncRNA *Airn* induces gene silencing and recruits Polycomb Repressive Complexes (PRCs) to varying extents over a 15 megabase domain. The mechanisms remain unclear. Using high-resolution approaches, we show in mouse trophoblast stem cells that *Airn* expression induces long-range changes to chromatin architecture that coincide with PRC-directed modifications and center around CpG island promoters that contact the *Airn* locus even in the absence of *Airn* expression. Intensity of contact between *Airn* lncRNA and target chromatin correlated with underlying intensity of PRC-directed chromatin modifications. Deletion of CpG islands that form contacts with *Airn* altered long-distance silencing and PRC activity in a manner that correlated with changes in chromatin architecture. We conclude that *Airn* is a potent *cis*-acting lncRNA whose primary functions of transcriptional repression and PRC recruitment are controlled by an equilibratory network of DNA regulatory elements that modulate its frequency of contact with target chromatin.

## INTRODUCTION

Genomic imprinting is a process known to occur for some ∼150 mammalian genes, resulting in their preferential expression from one parentally inherited allele over the other. Dysregulated imprinting can lead to developmental disorders, cancer, and changes in metabolism. Studies of genomic imprinting have also yielded other important insights, including the recognition that long noncoding RNAs (lncRNAs) control the expression of genes that are essential for proper human development (Monk et al. 2019; Tucci et al. 2019; MacDonald and Mann 2020; Lleres et al. 2021).

*Airn* (*<u>A</u>ntisense of Igf2r Non-Protein Coding <u>RN</u>A*) is a gene that in mice is imprinted and produces a lncRNA specifically from the paternal allele of chromosome 17. The *Airn* locus produces lncRNA transcripts that are upwards of ∼90kb in length but have heterogeneous 3′ ends (Huang et al. 2011). In addition, *Airn* transcripts are lowly abundant, short-lived, retained near their site of transcription, predominantly unspliced, and not conserved outside of rodents (Lyle et al. 2000; Killian et al. 2001; Sleutels et al. 2002; Seidl et al. 2006; Terranova et al. 2008; Yotova et al. 2008; Suzuki et al. 2018). Nevertheless, *Airn* expression results in transcriptional repression over a domain that spans ∼15 megabases (Mb) in extraembryonic tissues of the mouse, the largest autosomal region known to be under the control of a repressive lncRNA (Andergassen et al. 2017; Schertzer et al. 2019a). Repression by *Airn* occurs predominantly if not exclusively *in cis*, on the same chromosome from which the lncRNA is transcribed (Andergassen et al. 2017; Schertzer et al. 2019a).

The mechanism by which *Airn* induces repression over its 15Mb target domain is not clear. *Airn*’s lack of conservation, lack of splicing, and the instability and variable length of its RNA product have raised questions about whether it is the *Airn* lncRNA itself or merely the act of its transcription that induces repression (Latos et al. 2012; Pauler et al. 2012). Accumulating data are supportive of a role for the *Airn* lncRNA product in inducing long-range repression (Nagano et al. 2008; Andergassen et al. 2019; Schertzer et al. 2019a), yet it remains unclear what properties of the RNA may enable it to do so. Moreover, the intensity of repression across the *Airn* target domain is non-uniform (Schertzer et al. 2019a), implying that features of the genome influence repression by *Airn* in ways that are not yet clear.

For its full repressive effect, *Airn* requires several histone-modifying enzymes, including the Polycomb Repressive Complexes (PRCs), which are known to be recruited to chromatin by the expression of a handful of other lncRNAs, including *Xist* during the process of X chromosome Inactivation (Grosswendt et al. 2020; Andergassen et al. 2021; Trotman et al. 2021)). There are two major PRCs, PRC1 and PRC2, each of which can be classified into different sub-complexes that contain core and auxiliary components (Blackledge and Klose 2021). PRC1 monoubiquitinates histone H2A at lysine 119 (H2AK119ub) and is comprised of canonical and variant complexes termed cPRC1 and vPRC1, respectively. PRC2 tri-methylates histone H3 at lysine 27 (H3K27me3) and is comprised of sub-complexes called PRC2.1 and PRC2.2. The different auxiliary factors that distinguish PRC sub-complexes modulate their enzymatic activities, interaction partners, effects on 3-dimensional (3D) genome organization, and ultimately, effects on gene expression (Blackledge and Klose 2021). For example, specific forms of vPRC1 have been shown to interface the most closely with the lncRNA *Xist* (Almeida et al. 2017; Bousard et al. 2019).

We previously found that in mouse trophoblast stem cells (TSCs), expression of *Airn* induces gene silencing and the deposition of H2AK119ub and H3K27me3 in a highly non-uniform fashion across a 15Mb target domain (Schertzer et al. 2019a). Intensity of gene repression and PRC-directed modifications could be modulated by altering levels of *Airn* transcription from its endogenous promoter, supporting a role for the *Airn* lncRNA product in PRC recruitment and indicating that chromatin within its target domain is highly sensitive to the levels of *Airn*. Within the domain, the regions most highly decorated in PRC-deposited modifications centered around a subset of CpG island (CGI) promoters bound by the catalytic components of PRC1 and PRC2, even on the maternal allele, which does not express *Airn*. These and other data led us to hypothesize that the non-uniform repression across the *Airn* target domain was mediated by DNA regulatory elements that preferentially contact the *Airn* locus through 3D space and focus *Airn*‘s repressive activity over certain genomic regions.

Herein, we set out to test that hypothesis and examine in greater detail the extent to which *Airn*-induced silencing is influenced by chromatin architecture and underlying features of the genome. Indeed, in TSCs and mouse embryonic stem cells (ESCs), we found that variation in silencing across the domain could be partly explained by 3D DNA contacts that exist in the absence of *Airn* expression, which serve to bring certain genomic regions in closer proximity to the *Airn* locus over others. Regions within the domain that were silenced the most intensely were the ones that associated the most robustly with the *Airn* lncRNA product. Seemingly similar DNA regulatory elements located in different regions within the target domain had different effects on *Airn*-induced silencing, likely owing to roles in modulating local PRC recruitment and frequency of contact between *Airn* and target DNA. Our data support the notion that primary functions of the *Airn* lncRNA product are to repress gene expression and recruit multiple forms of PRC1 and PRC2 to proximal regions of chromatin. Furthermore, our work illustrates multiple examples of how DNA regulatory elements can control the regional intensity and genomic range of repression induced by a mammalian lncRNA, unveiling principles that likely apply elsewhere in the genome, in particular, to other domains that surround strong locus control regions.

## RESULTS

### *Airn* expression induces large-scale changes to chromatin architecture

Because *Airn* is monoallelically expressed and *cis*-acting, its target domain exists in different states on each allele; the paternal allele being repressed by *Airn*, and the maternal allele existing in the non-repressed state. For this reason, we and others have studied *Airn* in F1-hybrid cell lines or animals derived from different strains of inbred mice (Andergassen et al. 2017; Andergassen et al. 2019; Schertzer et al. 2019a). In F1-hybrids, physical events associated with maternal and paternal alleles can be distinguished by single-nucleotide polymorphisms (SNPs) in sequencing reads (Keane et al. 2011). The ability to simultaneously monitor repressed and non-repressed alleles in F1-hybrids makes them highly controlled settings in which to study monoallelically expressed lncRNAs such as *Airn*.

To determine how *Airn* expression alters chromatin architecture, we performed *in situ* Hi-C (Rao et al. 2014) in three F1-hybrid TSC lines: one line derived from a cross between a CAST/EiJ mother and C57BL/6J father (C/B TSCs), one derived from the reciprocal cross, a C57BL/6J mother and CAST/EiJ father (B/C TSCs), and a third TSC line in which we previously used CRISPR to insert a triple-polyadenylation cassette ∼3kb downstream of the *Airn* transcription start site in C/B TSCs, thereby generating a mutant with an expected null phenotype (C/B *Airn* truncation TSCs; (Sleutels et al. 2002; Schertzer et al. 2019a)). Hi-C libraries were generated in biological triplicate from each TSC line and sequenced to an aggregate depth per genotype of at least 1.6 billion paired-end 150 nucleotide (nt) reads. The Hi-C data across replicates and genotypes were reproducible and of high quality (Table S1).

We used Juicer and the mm9 genome to construct allele-specific contact maps in each TSC line, and first examined in broad strokes how *Airn* expression alters chromatin architecture across its target domain (Durand et al. 2016a; Durand et al. 2016b). In C/B WT TSCs, on the paternal B6 allele, which expresses *Airn*, relative to the maternal CAST allele, which does not, we observed a reduction in short-range DNA contacts concomitant with an increase in long-range contacts, which were largely contained within the 15Mb target domain previously shown to be repressed by *Airn* in TSC (Figures 1A and B, panel (i); (Schertzer et al. 2019a)). The greatest increases in contact occurred within a 4.5Mb interval that extended from the *Airn* locus and terminated at the genes *Prr18*, *T*, and *Pde10a* (Figures 1A and B, panel (i)). In *Airn* truncation TSCs, the increased contacts were not detectable, demonstrating their dependence on *Airn* expression (Figures 1A and B, panel (ii)). Moreover, the overall trends that were observed in C/B WT TSCs were also observed in the reciprocal F1-hybrid wildtype cell line -- B/C TSCs -- in which the paternal, *Airn*- expressing allele is of CAST origin, confirming that differences in chromatin architecture between paternal and maternal alleles are due to parent-of-origin and not strain-specific effects (Figures 1A and B, panel (iii)). The regions that underwent the strongest *Airn*-induced changes in contact frequency were the ones that most clearly shifted from the “A” to the “B” chromosome compartment specifically on the alleles that expressed *Airn* (Figure 1C, panels (i-iii); Figure S1; (Lieberman-Aiden et al. 2009; Rao et al. 2014)). These data demonstrate that in TSCs, *Airn* expression induces large-scale changes to chromatin architecture that are largely contained within a 15Mb genomic interval previously shown to be subject to *Airn*-dependent repression (Andergassen et al. 2017; Schertzer et al. 2019a).

**Figure 1.**
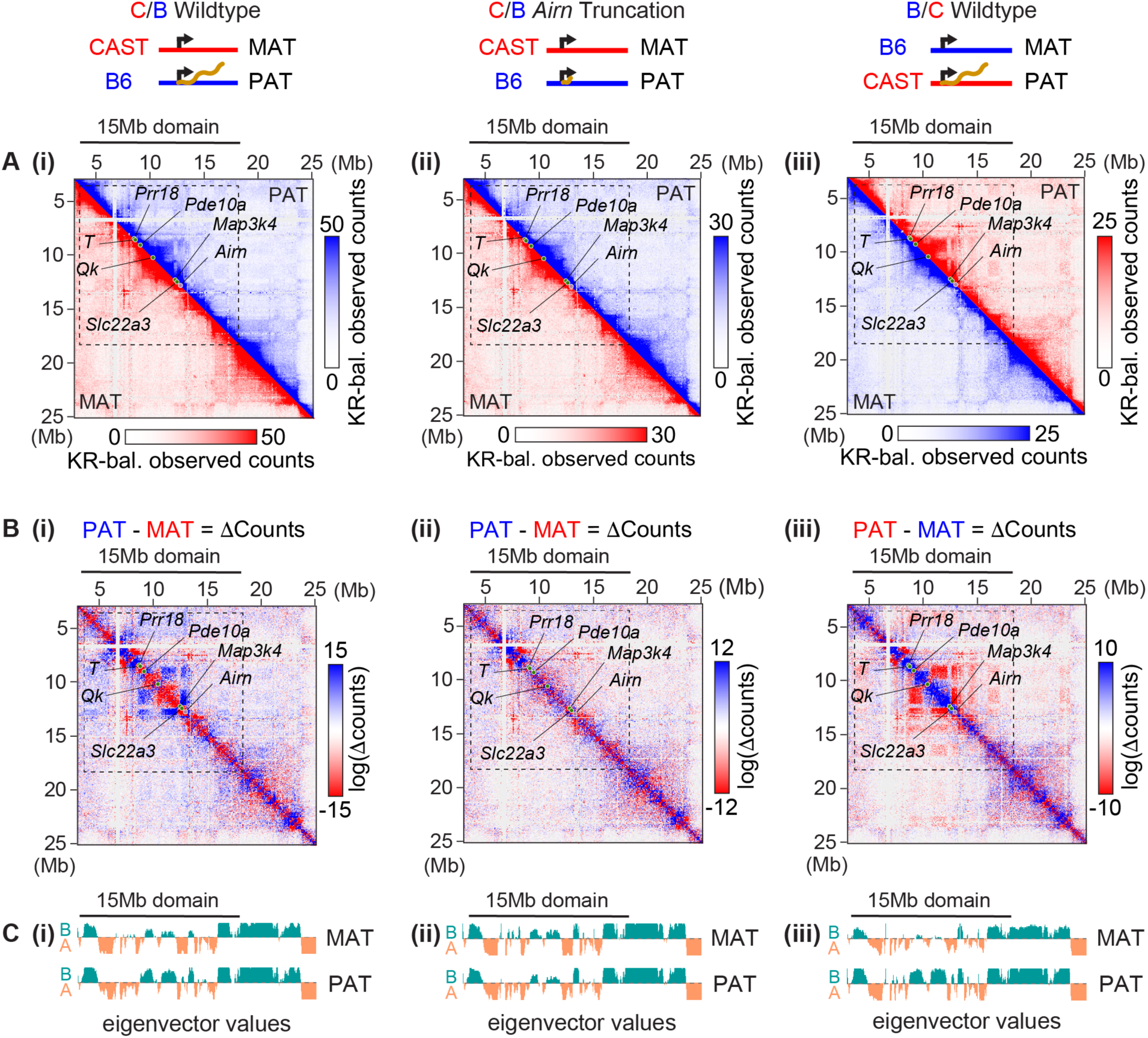
*Airn* expression induces large-scale changes to chromatin architecture. **(A)** 2D Hi-C contact heatmaps of allelic observed counts in **(i)** C/B wildtype, **(ii)** C/B *Airn* truncation, and **(iii)** B/C wildtype TSCs. Maternal (bottom left) and paternal (top right) heatmaps are partitioned, KR-balanced, and 50kb resolution. KR bal., Knight-Ruiz balanced. **(B)** Subtraction contact heatmaps of [PAT minus MAT] observed counts, (i-iii) as (A)**. (C)** Density tracks of eigenvector values at 50kb resolution for “A” and “B” chromosome compartmentalization. In all heatmaps: dotted lines, 15Mb *Airn* target domain; purple circle, *Airn* gene; green circles, other loci of interest.

### *Airn*-induced changes in chromatin architecture coincide with the presence of PRC- deposited modifications

To gain a better sense of how DNA contacts with the *Airn* locus correlate with *Airn*-dependent repression, we created a series of “viewpoint” plots, in which contacts between *Airn* and surrounding regions were extracted and visualized in two dimensions. Consistent with the heatmaps of Figure 1, we observed a dramatic increase in *Airn*-induced contacts with the *Airn* locus on the paternal allele towards the centromeric end of chr17, including a pronounced shoulder of increased contacts surrounding the genes *Prr18*, *T*, and *Pde10a* (Figure 2A). We also observed augmented contacts between *Airn* and the gene *Qk* on both alleles in all TSC lines profiled (Figure 2A). In C/B TSCs, the intensity of allele-specific contacts with *Airn* as measured by Hi-C correlated remarkably well with allele-specific distances to the *Airn* locus that we had previously measured by DNA FISH, corroborating both forms of measurement (Figure 2B; Spearman’s ρ of -0.82 and -0.95 and p = 0.007 and 0.001 on maternal and paternal alleles, respectively; FISH probe locations from (Schertzer et al. 2019a) shown under panel A(i)).

**Figure 2.**
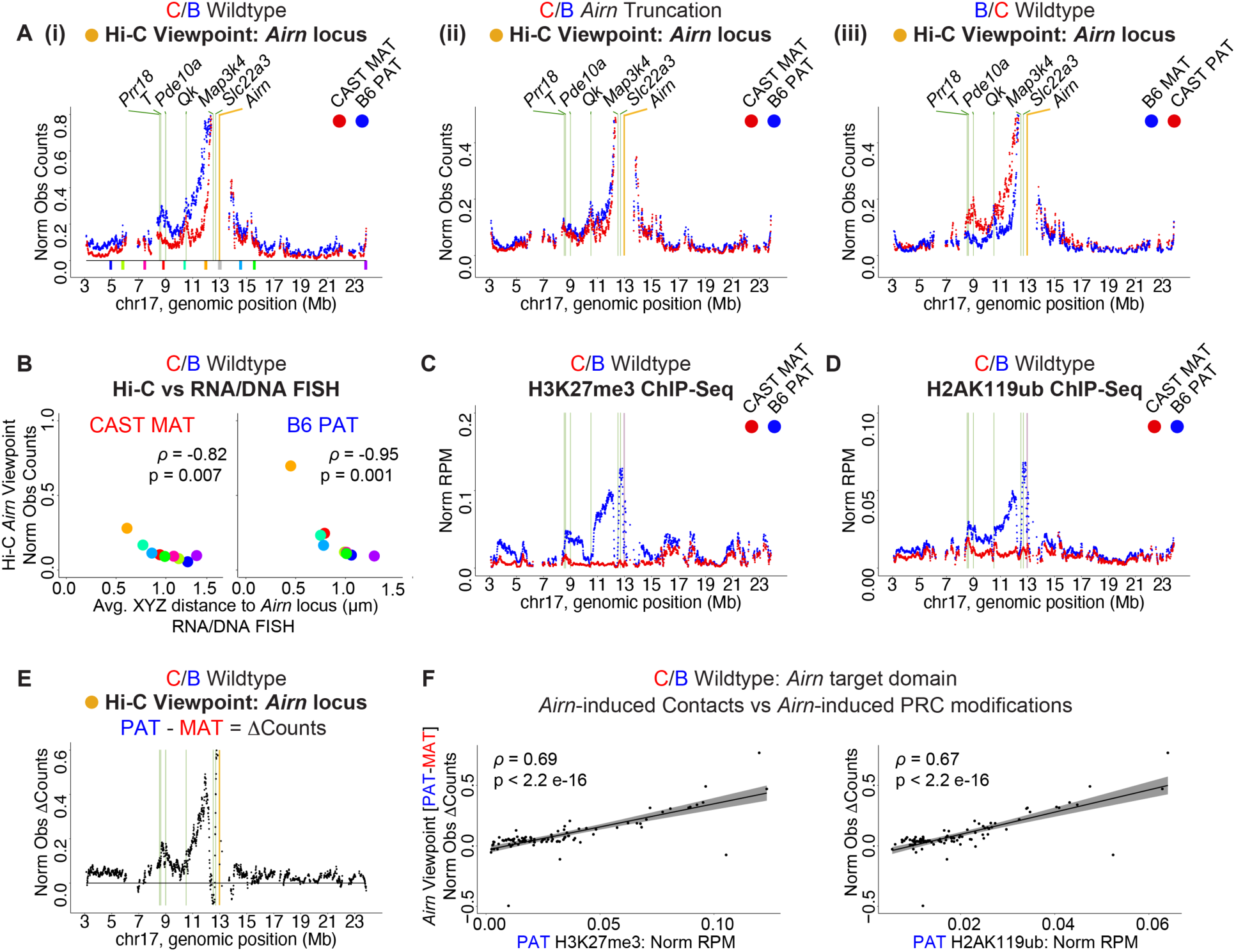
*Airn*-induced changes in chromatin architecture coincide with the presence of PRC-deposited modifications. **(A)** Tiling density plots of allelic Hi-C *Airn* viewpoint observed contact counts in **(i)** C/B wildtype, **(ii)** C/B *Airn* truncation, and **(iii)** B/C wildtype TSCs. Colored blocks in (i), FISH probes analyzed in (B; (Schertzer et al. 2019a)). **(B)** Allelic Hi-C *Airn* viewpoint contacts from (A, panel (i)) vs average distance to *Airn* measured by RNA/DNA FISH in C/B wildtype TSCs from (Schertzer et al. 2019a). Spearman’s ρ and p values are shown. **(C)** Tiling density plot of allelic H3K27me3 ChIP-Seq data in C/B wildtype TSCs from (Schertzer et al. 2019a). **(D)** Tiling density plot of allelic H2AK119ub ChIP-Seq data in C/B wildtype TSCs from (Schertzer et al. 2019a). **(E)** Tiling density plot of allelic *Airn* viewpoint [PAT minus MAT] observed counts in C/B wildtype TSCs from (A, panel (i)). **(F)** Scatter plots of *Airn* viewpoint [PAT minus MAT] observed counts vs H3K27me3 (left) and H2AK119ub (right). Spearman’s ρ and p values are shown. In all tiling density plots, counts were summed in 10kb bins, normalized for SNP density, then averaged across 9bin windows in 1bin tiling increments. For viewpoint plots, we excluded bins whose aggregate SNP-overlapping read count across merged Hi-C datasets fell within the bottom quintile relative to the rest of the genome. For ChIP-Seq plots, bins with greater or equal to 25 B6/CAST SNPs are shown. RPM, Reads per Million total reads. In all panels: yellow bar, viewpoint; purple bar, *Airn* gene; green bars, other loci of interest.

The magnitude of *Airn*-induced contacts (i.e., paternal contacts subtracted from maternal contacts) correlated remarkably well with the underlying intensity of *Airn*-dependent, PRC-directed chromatin modifications (Figures 2C-F; Spearman’s ρ between *Airn*-induced contacts and H3K27me3 and H2AK119ub in C/B TSCs, 0.69 and 0.67, respectively, p < 2e-16 for both comparisons). Moreover, we used ChIP-Seq to profile seven individual components of PRC1 and PRC2, and six out of the seven components exhibited enrichment within the target domain, specifically on the *Airn*-expressing paternal allele (Figure S2A). The sole exception was the vPRC1 component KDM2B, which did not exhibit evidence of responsiveness to *Airn* or to the lncRNA *Xist* (Figures S2A and B). These data, alongside prior data indicating that the PRCs and their modifications to chromatin can induce DNA compaction (Terranova et al. 2008; Rao et al. 2014; Blackledge and Klose 2021), are consistent with the notion that the large-scale changes in genome architecture induced by the expression of *Airn* depend, at least in part, on the PRCs and their modifications to chromatin.

### *Airn*-induced repression centers around regions that form pre-existing contacts with the *Airn* locus and harbor CGIs bound by vPRC1

DNA contacts detected by Hi-C can occur by chance with a frequency that increases proportionally with decreasing distance from the locus in question (Rao et al. 2014). To gain a sense of the relative intensity of contacts that occur with the *Airn* locus after correcting for distance-dependent effects, we created a series of Observed-over-Expected (O/E) plots with *Airn* as the viewpoint, in which detected contacts were normalized by those expected from a distance-dependent decay model (Rao et al. 2014).

These O/E viewpoint plots revealed three local maxima of contact with *Airn* that fell within the 4.5Mb genomic interval that is the most intensely repressed by *Airn*, extending from the *Airn* locus and terminating at *Prr18*, *T*, and *Pde10a* (Figures 2 and 3A, panels (i-iii)). Specifically, maxima were detected surrounding the genes *Prr18*/*T*/*Pde10a*, the gene *Qk*, and the gene *Slc22a3* (Figure 3A, panels (i-iii)). While the intensity of O/E contact with *Prr18*/*T*/*Pde10a* increased dramatically upon expression of *Airn*, the intensity of O/E contact with *Qk* and *Slc22a3* changed to lesser extents or not at all (Figure 3A, panels (i-iii)). Importantly, all three maxima were clearly present even in *Airn* truncation TSCs, highlighting their ability to contact the *Airn* locus even in the absence of *Airn* expression (Figure 3A, panel (ii)). Of note, each of the genes within these maxima are driven by CGI promoters that we found in previous work either bind high levels of RING1B and EZH2 (in the cases of *Prr18*/*T*/*Pde10a* and *Qk*), or are present in the region of the *Airn* target domain that accumulates the highest levels of *Airn*-induced, PRC-directed chromatin modifications (in the case of *Slc22a3*; Figure 2; (Schertzer et al. 2019a)).

**Figure 3.**
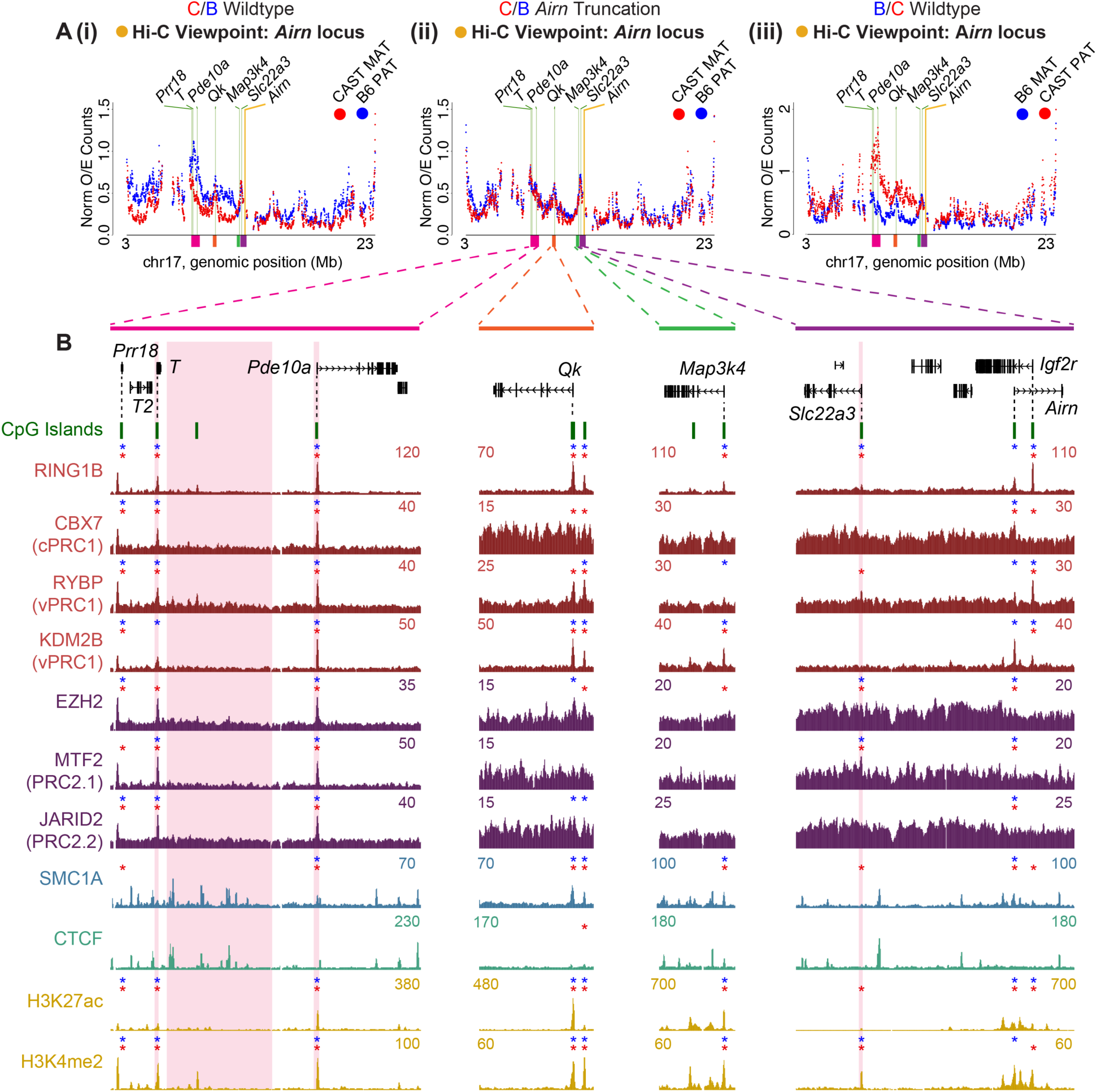
***Airn*-induced repression centers around regions that form pre-existing contacts with the *Airn* locus and harbor CGIs bound by vPRC1.** (A) Tiling density plots of allelic Hi-C *Airn* viewpoint Observed-over-Expected (O/E) counts in (i) C/B wildtype, (ii) C/B *Airn* truncation, and (iii) B/C wildtype TSCs. Data processed as in Figure 2. In all plots: yellow bar, viewpoint; green bars, other loci of interest. (B) Genome Browser graphics of the *Airn* contact domain, *Map3k4*, *Qk*, and *Prr18/T/Pde10a*. ChIP-Seq tracks display non-allelic read density. Red or blue asterisks, CGIs exhibiting significant enrichment of the factor on the maternal or paternal alleles, respectively (p<0.05, permutation). See Table S3. Pink rectangles, DNA regions deleted in Figures 5-7.

For several reasons, we were intrigued that points of contact with *Airn* were centered around PRC-bound CGIs. In both ESCs and TSCs, CGIs often mark high-density sites of PRC binding, and in many cell types, PRCs and the modifications that they deposit on chromatin have been shown to mediate long-range 3D contacts independently of the major architectural factors CTCF and Cohesin (Isono et al. 2013; Oksuz et al. 2018; Cuadrado et al. 2019; Blackledge et al. 2020; Boyle et al. 2020; Rhodes et al. 2020; Cai et al. 2021; Eeftens et al. 2021; Kriz et al. 2021; Kraft et al. 2022). Moreover, the genes within the points of contact -- *Pde10a*, *Qk*, and *Slc22a3* -- are all repressed by *Airn* in TSCs ((Schertzer et al. 2019a); Table S2). Also, the *Airn* gene body itself harbors two CGIs that bind RING1B/PRC1 (Schertzer et al. 2019a), and the *Airn* lncRNA has previously been found to associate with the *Slc22a3* CGI (Nagano et al. 2008). Lastly, although the reasons remain unclear, we previously found that deletion of the *Slc22a3* CGI resulted in a dramatic loss of *Airn*-induced accumulation of H3K27me3, most notably in the 4.5Mb interval beginning at *Airn* and terminating at *Prr18*, *T*, and *Pde10a* (Schertzer et al. 2019a). All together, these data raise the possibility that features associated with the CGI promoters found in regions that form augmented contacts with *Airn* play roles in modulating the local intensity of *Airn*-induced repression.

With the above possibility in mind, we used ChIP-Seq to examine what factors and chromatin modifications were enriched over CGIs contained within points of 3D contact with *Airn*. These included CGIs at *Pde10a*, *Qk*, *Slc22a3*, and *Airn* itself, as well as the CGI promoter of the gene *Map3k4*. Like *Qk*, *Map3k4* is repressed by *Airn* and forms a detectable contact with the *Airn* locus by Hi-C (Figure S3), but partially escapes silencing and sits within a region that resists the local accumulation of *Airn*-induced, PRC-deposited chromatin modifications (Figures 2C and D; Table S2).

In total, we examined ChIP-Seq data for seven individual PRC components (RING1B, RYBP, CBX7, KDM2B, EZH2, MTF2, and JARID2), four chromatin modifications (H3K27me3, H2K119ub, H3K4me2, and H3K27ac), and two architectural factors (SMC1A/Cohesin and CTCF; data from this study and (Calabrese et al. 2012; Schertzer et al. 2019a)). A summary of the allele-specific enrichment of each factor is found in Table S3, and the non-allelic UCSC wiggle density tracks are shown in (Figure 3B). The asterisks above each CGI indicate whether the factor was detected on the maternal allele, paternal allele, or both (Figure 3B; Table S3).

While there was not one singular pattern of enrichment, notable similarities did emerge. Each CGI except for the one found at the promoter of *Airn* showed some level of peak-like enrichment for vPRC1 on the maternal allele (Figure 3B). Likewise, all of the CGIs examined except for the one at the *Airn* promoter showed peak-like enrichment of at least one of the two chromatin modifications associated with transcriptional activation (H3K4me2 or H3K27ac), also on the maternal allele. SMC1A/Cohesin was also detected on the maternal allele of these same set of CGIs, although its intensity of enrichment was low relative to intergenic peaks (Figure 3B). Thus, the regions within the target domain that form contacts with the *Airn* locus on the maternal allele all harbor CGI promoters that are enriched in vPRC1, Cohesin, and signatures of transcriptional activity.

In contrast, CGIs in the region that underwent the strongest *Airn*-induced changes in chromatin architecture, surrounding the genes *Prr18*, *T*, and *Pde10a*, were associated with sharp peaks of cPRC1 and PRC2 as well as vPRC1 (Figures 3B and S2A). Of those CGIs, the CGI at *Pde10a* associated with the highest levels of both PRC1 and PRC2 (Table S3). Likewise, the intergenic regions that underwent the strongest *Airn*-induced changes in chromatin architecture were similarly enriched in cPRC1 and PRC2 as well as vPRC1 (Figure S2A). Thus, while the presence of vPRC1, Cohesin, and transcription can identify regions that contact the *Airn* locus in the absence of *Airn* expression, the presence of cPRC1 and PRC2 at both CGIs and intergenic regions correlate more strongly with the presence of *Airn*-induced, PRC-deposited modifications and long-range changes to architecture.

### Presence of *Airn* lncRNA on chromatin correlates with PRC-deposited modifications and centers around pre-existing contacts with the *Airn* locus

Considering that across the *Airn* target domain, we observed variations in the intensity of DNA contacts, associated proteins, PRC-directed modifications, and gene silencing, we sought to determine whether the *Airn* lncRNA itself preferentially associated with specific DNA regions or regulatory elements. To address this question, we used CHART-Seq, an approach to identify genomic regions located in proximity to lncRNAs of interest (Simon et al. 2013). Utilizing a set of 51 DNA oligos spaced across the first 75kb of the *Airn* lncRNA (Figure S4A), we performed CHART-Seq in C/B wildtype TSCs, in *Airn* truncation TSCs, and in a C/B TSC line from (Schertzer et al. 2019a), in which we used CRISPR-Cas9 to over-express *Airn* from its endogenous promoter (*Airn* Highly-Expressing or H-E TSCs). RT-qPCR confirmed expected levels of *Airn* expression in each TSC line (Figure S4B).

In C/B wildtype TSCs, *Airn* CHART-Seq revealed enrichment of DNA on the paternal but not maternal allele across the *Airn* target domain, beginning near the centromere and ending ∼3Mb downstream of the *Airn* locus, the same region where the last *Airn*-induced PRC-dependent modifications are visible (Figures 4A and B, panels (i)). In H-E TSCs, the enrichment increased commensurately with the ∼3-fold over-expression of *Airn*, including an expanded range over which associations with *Airn* were detected (Figures 4A, panel (ii)). Conversely, in *Airn* truncation cells, the enrichment of DNA from the paternal allele was lost (Figure 4A, panel (iii)). Together, these data indicate that the DNA recovered by CHART depends on and is sensitive to the overall levels of *Airn* expression. Moreover, we observed a remarkably strong correlation between *Airn* CHART-Seq and H3K27me3 ChIP-Seq signal on the paternal allele throughout the target domain (Spearman’s ρ in C/B wildtype and H-E TSCs, 0.63 and 0.74, respectively; p < 2.2 e-16 in each comparison; Figure 4A vs 4B). We were especially struck by the lower CHART-Seq signal in H-E TSCs that began just upstream of *Map3k4* and *Qk*, genes that are repressed by *Airn* but located at inflection points where the intensity of PRC-directed modifications drops precipitously (Figures 4A and B). Likewise, particularly in H-E TSCs, local maxima of *Airn* association coincide with genomic regions that form local maxima of DNA contacts with the *Airn* locus even in the absence of *Airn* expression (Figure 4C). Thus, these data support the notions that a major function of the *Airn* lncRNA product is to recruit the PRCs to modify chromatin over a 15Mb domain, and that proximity to *Airn* lncRNA dictates the intensity with which these modifications occur. Moreover, pre-existing DNA contacts – those that occur with the *Airn* locus in the absence of *Airn* expression – identify which regions of chromatin will contact the *Airn* lncRNA and become decorated in PRC-directed modifications. Lastly, features located within or near the CGI promoters of *Map3k4* and *Qk* may enable those genes to resist forming contacts with the *Airn* lncRNA that are detectable by CHART.

**Figure 4.**
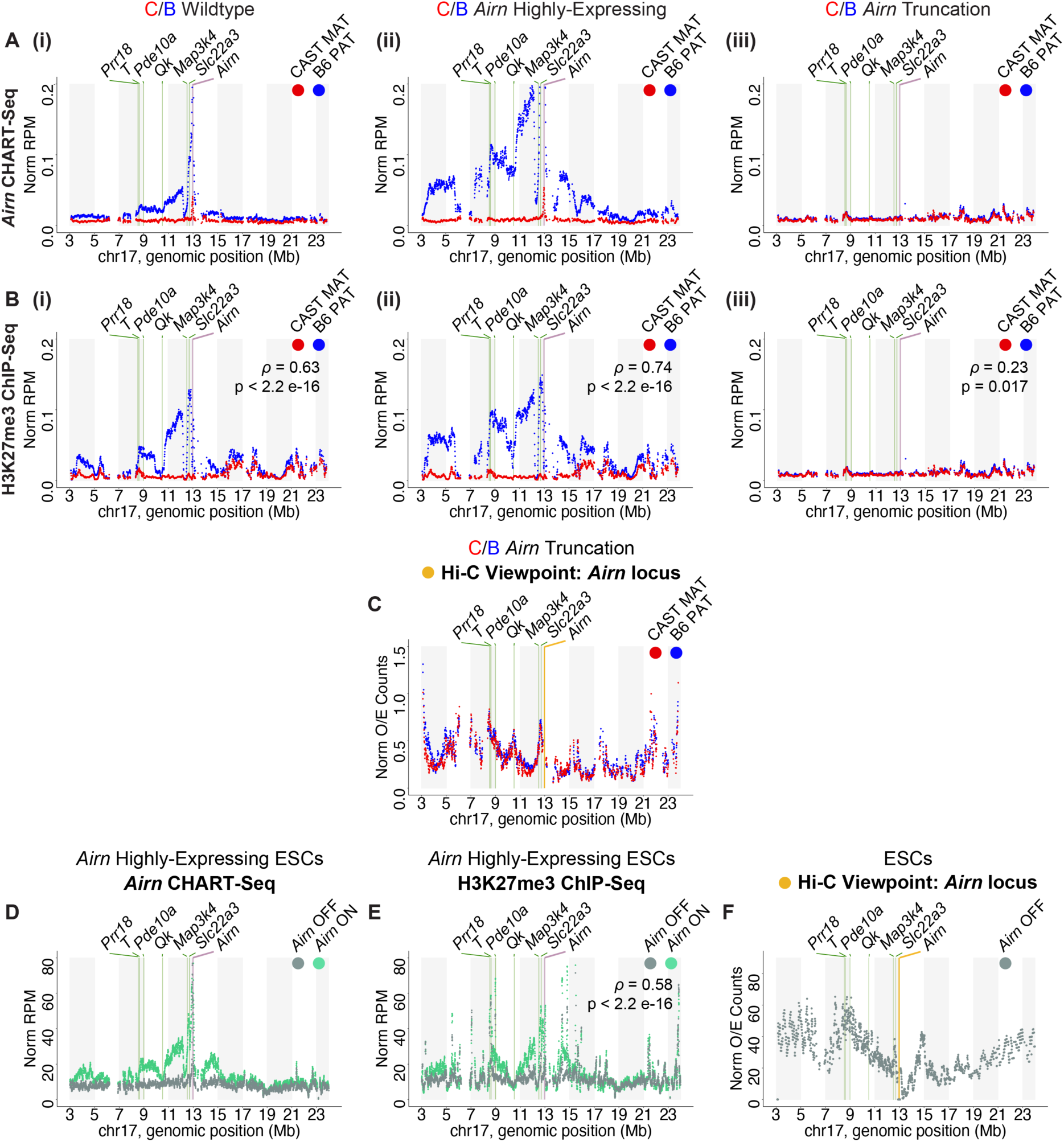
Presence of *Airn* lncRNA on chromatin correlates with PRC-deposited modifications and centers around pre-existing contacts with the *Airn* locus. (A) Tiling density plots of allelic *Airn* CHART-Seq data in (i) C/B wildtype, (ii) C/B *Airn* High-expressing (H-E), and (iii) C/B *Airn* truncation TSCs. (B) Tiling density plots of allelic H3K27me3 ChIP-Seq data from (Schertzer et al. 2019a), (i-iii) as (A). (C) Tiling density plots of allelic Hi-C *Airn* viewpoint Observed-over-Expected (O/E) counts in C/B *Airn* truncation TSCs. (D) Tiling density plots of non-allelic *Airn* CHART-Seq data from untreated (*Airn* OFF) or dox. treated H-E ESCs (*Airn* ON). (E) Tiling density plots of non-allelic H3K27me3 ChIP-Seq data. (F) Tiling density plot of non-allelic Hi-C *Airn* viewpoint O/E counts in ESCs (Dixon et al. 2012). All tiling density plots but those in (D) and (E) were generated as in Figure 2. Read counts in (D) and (F) were summed in 40kb bins tiled in 4kb increments across the chromosome, converted to RPM (Reads per Million total reads), and then normalized and filtered for at least 50% genome read alignability (see Methods). In all panels: purple bar, *Airn* gene; green bars, other loci of interest.

In TSCs and the extraembryonic lineages, *Airn* is expressed, represses genes, and recruits PRCs over a 15Mb genomic domain (Andergassen et al. 2017; Schertzer et al. 2019a). However, in ESCs, *Airn* is not expressed (Latos et al. 2009). If gene repression and PRC recruitment is an inherent function of the *Airn* lncRNA during the earliest stages of mouse embryogenesis, then we might expect its forced expression in ESCs to result in the accumulation of the *Airn* lncRNA and PRC-deposited modifications over a target domain similar in size to the one found in TSCs. If, in contrast, the function of *Airn* is somehow restricted to the extraembryonic lineages, its forced expression in ESCs would have little effect.

To test these hypotheses, we performed *Airn* CHART-Seq in mouse ESCs in which we used CRISPR-Cas9 technology to force expression of *Airn* from its own promoter (H-E ESCs). In H-E ESCs, *Airn* was expressed at a level approximately equal to WT TSCs (Figure S4B). As a negative control, we also performed CHART-Seq in uninduced ESCs. In parallel, we performed H3K27me3 ChIP-Seq in these same ESCs to determine the extent to which DNA retrieved by CHART correlated with intensity of H3K27me3. Consistent with *Airn* being capable of inducing repression even in ESCs, we observed associations between the *Airn* lncRNA and DNA in H-E ESCs across a domain that was remarkably similar in size and contour to the domain contacted by *Airn* in TSCs (Figure 4D vs 4A). As in TSCs, *Airn* contacts were significantly correlated with underlying H3K27me3 (Figure 4D vs 4E; Spearman’s ρ = 0.58). Moreover, again similar to TSCs, *Map3k4* and *Qk* resisted contacts with *Airn* and PRC-directed modifications in H-E ESCs (Figures 4D and E). Analyzing previously published ESC Hi-C data (Dixon et al. 2012), we found that Observed-over-Expected contacts with the *Airn* locus essentially predict which regions of DNA will contact *Airn* lncRNA and accumulate H3K27me3 (Figure 4F vs 4D, E). Thus, even in ESCs, the *Airn* lncRNA is capable of recruiting PRCs throughout a broad domain, and its effects are influenced by pre-existing structure of the genome.

### DNA regulatory element deletions alter levels of PRC-directed modifications and gene silencing throughout the *Airn* target domain

Our data indicate that regional proximity to the *Airn* lncRNA product is linked to the local intensity of gene silencing and recruitment of PRCs. To better understand the roles that specific DNA regulatory elements might play in controlling proximity to *Airn*, we focused on the region surrounding the genes *Prr18*/*T*/*Pde10a*, which contains several CGIs that robustly bind the PRCs and undergoes increased frequency of contact with the *Airn* locus upon *Airn* expression (Figures 2 and 3). We used CRISPR to individually delete the CGI promoters of *T* and *Pde10a*, as well as a 190kb cluster of intergenic CTCF and SMC1A/Cohesin peaks located between *T* and *Pde10a* (pink rectangles in Figure 3B; Figures S5A-C). We were able to derive heterozygous clonal TSC lines harboring deletions for each element on their paternal alleles (Δ*T*, four lines; Δ*Pde10a*, two lines; Δ*Cluster*, two lines; Figures S5A-C). As controls, we derived four clonal non-targeting (NTG) TSC lines that harbor the same doxycycline-inducible *Cas9* transgene and underwent the same process of electroporation, clonal selection, and induction as above, but that express a non-targeting sgRNA that does not match the mouse genome. Additionally, we revived the two clonal TSC lines in which we previously deleted the *Slc22a3* CGI on the paternal allele (Δ*Slc22a3* TSCs from (Schertzer et al. 2019a); Figure S5D). In that previous study, we found that deletion of the *Slc22a3* CGI caused a ∼4.5Mb reduction in the intensity of H3K27me3, beginning essentially at the *Slc22a3* gene and extending through the distal cluster of PRC-bound CGIs at *Prr18*/*T*/*Pde10a* (Schertzer et al. 2019a).

Given that *Prr18*/*T*/*Pde10a* undergo pronounced increases in DNA contacts with both *Slc22a3* and *Airn*, specifically when *Airn* is expressed (Figures 2A and E; Figure S3A, D, E), we hypothesized that deletion of regulatory elements within the *Prr18*/*T*/*Pde10a* region would similarly disrupt DNA contacts and silencing by *Airn*, and also, that deletion of the *Slc22a3* CGI would likewise be accompanied by disruptions in long-range DNA contacts and silencing. RT-qPCR and RNA-Seq showed that *Airn* expression levels varied by no more than two-fold across our panel of lines (Figures S5E).

We performed H3K27me3 and H2AK119ub ChIP-Seq as well as RNA-Seq to examine how the deletions affected PRC activity and gene silencing. Relative to NTG controls, ChIP-Seq in Δ*Slc22a3* TSCs revealed a dramatic loss of H3K27me3 and H2AK119ub throughout the *Airn* target domain (Figure 5A vs S5F, G), both consistent with and extending results from (Schertzer et al. 2019a). However, whereas deletion of the *T* CGI had little to no effect, deletion of the *Pde10a* CGI unexpectedly caused a dramatic increase in the levels of H3K27me3 and H2AK119ub throughout the target domain, opposite to that observed in Δ*Slc22a3* TSCs (Figures 5B and C). Deletion of the cluster of CTCF and SMC1A/Cohesin peaks similarly increased H3K27me3 and H2AK119ub (Figure 5D). Cross-genotype comparisons within the target domain and across the remainder of chr17 are shown in Figure S5G.

**Figure 5.**
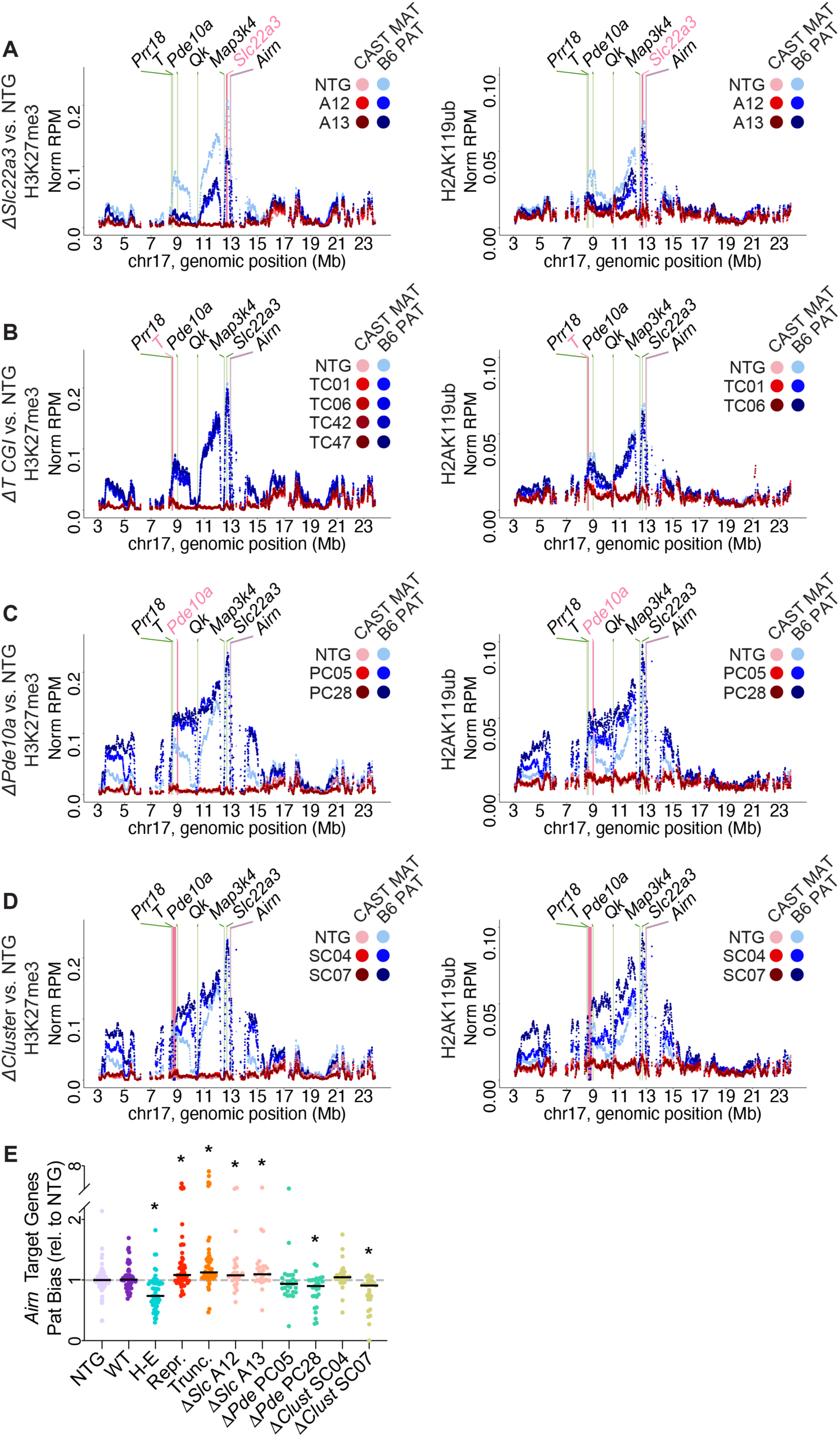
DNA regulatory element deletions alter levels of PRC-directed modifications and gene silencing throughout the *Airn* target domain. (A-D) Tiling density plots of allelic H3K27me3 (left) and H2AK119ub (right) ChIP-Seq data in (A) C/B Δ*Slc22a3*, (B) C/B *ΔT*, (C) C/B Δ*Pde10a* and (D) C/B Δ*Cluster* TSCs. Data from NTG clones were averaged and plotted in (A-D) and shown separately for comparison in Figure S5F. Data processed as in Fig. 2. (E) Proportion of paternal expression of the 27 *Airn* target genes relative to the averaged NTG value. Asterisks, significant changes relative to NTG (p<0.05, Welch t-test). In all tiling density plots: purple bar, *Airn* gene; green bars, other loci of interest; pink bars, DNA regions deleted.

RNA-Seq from deletion clones showed changes in gene expression consistent with changes in PRC-deposited modifications (Figure 5E vs A-D). In our previous study, we identified 27 genes within the target domain that were subject to repression by *Airn* (see Figures 4D and S5 from (Schertzer et al. 2019a)). In Δ*Slc22a3* TSCs, the relative paternal expression of these 27 genes increased significantly compared to their baseline in NTGs, up to an average level that was slightly less than that observed in *Airn* truncation TSCs, which are effectively null mutants ((Schertzer et al. 2019a); Figure 5E; NTG vs Δ*Slc22a3* (A12 and A13 clones), p = 0.038 and 0.025, respectively, Welch two sample t-test). Conversely, in Δ*Pde10a* and Δ*Cluster* TSCs, paternal expression of the 27 target genes decreased relative to NTG TSCs, albeit with variability between clones (Figure 5E; NTG vs Δ*Pde10a* and NTG vs Δ*Cluster*). The decreased paternal expression in Δ*Pde10a* and Δ*Cluster* TSCs was not as strong as that observed in *Airn* highly-expressing (H-E) TSCs (Figure 5E), but was nevertheless consistent with the increase in PRC-deposited modifications (Figures 5C and D; (Schertzer et al. 2019a)). Thus, seemingly similar DNA regulatory elements play critical and different roles in dictating the regional intensity of gene silencing and PRC recruitment induced by *Airn* within its15Mb target domain.

### Changes in DNA contacts with *Airn* mirror changes in PRC activity caused by regulatory element deletion

To gain mechanistic insight into the effects caused by regulatory element deletion, we used *in situ* Hi-C to examine DNA contacts in NTG, Δ*Slc22a3*, and Δ*Pde10a* TSCs. Hi-C data were of high quality and reproducible between clones (Table S1). Strikingly, consistent with changes in gene repression and PRC-deposited modifications, deletion of the *Slc22a3* and *Pde10a* CGIs alternately diminished and increased *Airn*-induced contacts across the entire target domain, coincident with corresponding changes in the intensities of compartmentalization (Figures 6A and B; Figure S6).

**Figure 6.**
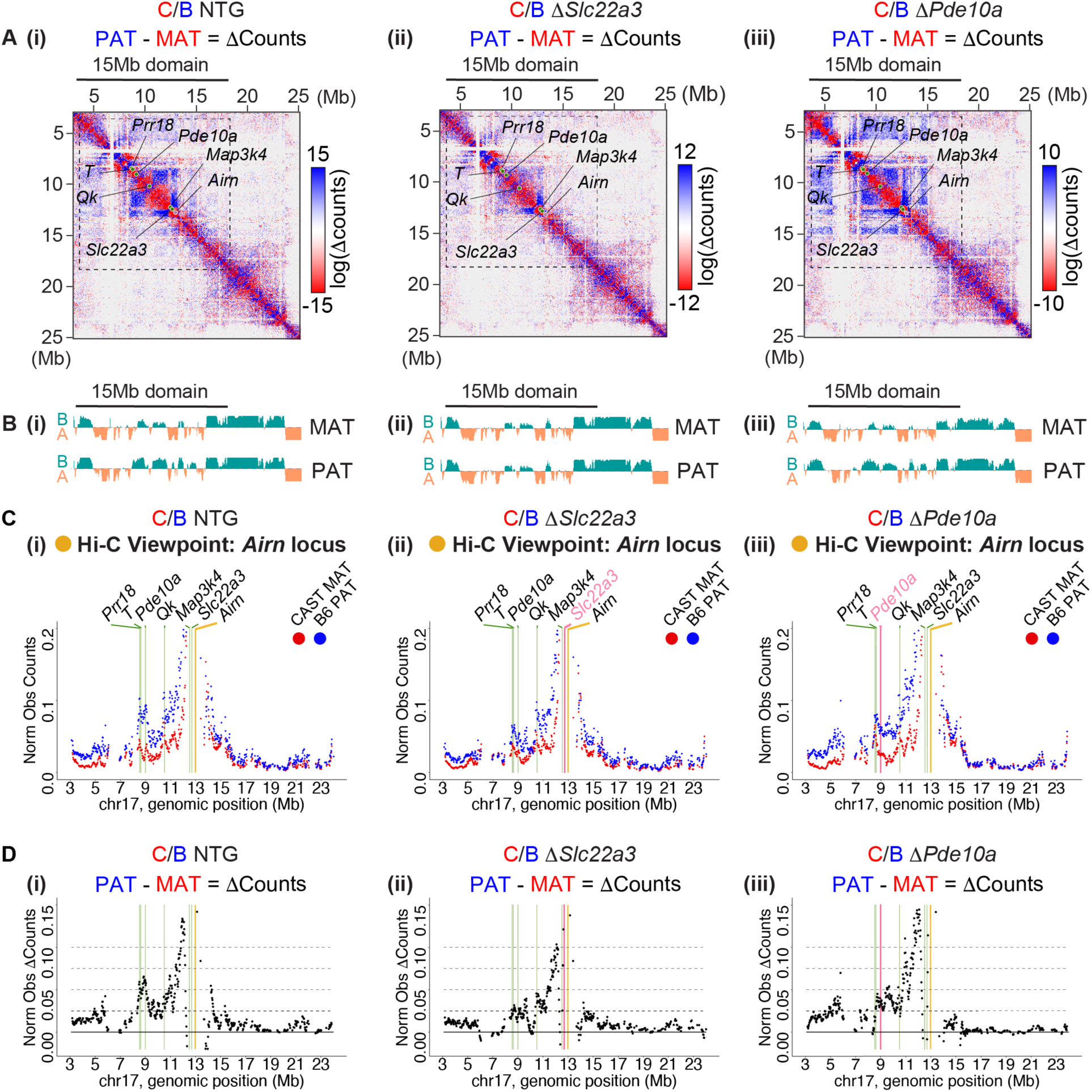
Changes in DNA contacts with *Airn* mirror changes in PRC activity caused by regulatory element deletion. **(A)** Hi-C subtraction contact heatmaps of [PAT minus MAT] observed counts in **(i)** C/B NTG, **(ii)** C/B Δ*Slc22a3*, and **(iii)** C/B Δ*Pde10a* TSCs. Counts are KR- balanced, 50kb resolution, and log2 transformed. **(B)** Density tracks of eigenvector values at 50kb resolution for “A” and “B” chromosome compartmentalization, (i-iii) as (A). **(C)** Tiling density plots of allelic *Airn* viewpoint observed contact counts, (i-iii) as (A). **(D)** Tiling density plots of allelic *Airn* viewpoint [PAT minus MAT] observed counts, (i-iii) as (A). All tiling density plots processed as in Figure 2. In all heatmaps: dotted lines, 15Mb *Airn* target domain; purple circles, *Airn* gene; green circles, other loci of interest. In all tiling density plots: yellow bar, *Airn* locus viewpoint; green bars, other loci of interest; pink bar, DNA region deleted.

Examining contacts from the *Airn* viewpoint provided additional insights (Figures 6C and D). Deletion of the *Slc22a3* CGI was coincident with reduced levels of *Airn*-induced contacts throughout the domain, with a possible exception at *Qk* (Figures 6C and D, panels (ii)). Proportionally, the greatest decreases in *Airn*-induced contacts surrounded *Prr18*/*T*/*Pde10a* (Figures 6C and D, panels (ii)). In contrast, in Δ*Pde10a* cells, *Airn*-induced contacts increased uniformly except at *Prr18*/*T*/*Pde10a* (Figures 6C and D, panels (iii)). Thus, deletion of the *Slc22a3* CGI reduced the interaction between *Airn* and DNA throughout its target domain, whereas deletion of the *Pde10a* CGI increased the interaction between *Airn* and all other regions in the domain save *Pde10a*.

### *Airn* expression is coincident with dissolution of DNA loops encasing *Slc22a3* and a local increase in PRC-directed modifications

We next considered a possible model whereby deletion of the *Slc22a3* CGI might restrict silencing by *Airn*. Examining our initial Hi-C datasets, we noted that the *Slc22a3* and *Airn* genes are located within the same contact domain, where they sit within nested DNA loops anchored by CTCF and Cohesin (Figures 7A and B). Moreover, by Hi-C, the loops that surround *Slc22a3* were reduced by *Airn* expression, to the extent that they are no longer detectable by the SIP algorithm on *Airn*-expressing alleles (Figure 7A; (Rowley et al. 2020)). Likewise, we observed a relative reduction in SMC1A and CTCF binding at those same loop anchors, again on *Airn*-expressing alleles, consistent with the reduced loop intensity (Figure 7B). Thus, it is conceivable that prior to *Airn* expression, the nested loops that encase *Slc22a3* and *Airn* may reduce the latter’s ability to interact with distal DNA. Upon *Airn* expression, recruitment of PRCs locally, promoted by the *Slc22a3* CGI, could antagonize Cohesin (Cuadrado et al. 2019; Rhodes et al. 2020; Kriz et al. 2021), disrupt loops, and enable *Airn* to contact distal regions of chromatin more efficiently.

**Figure 7.**
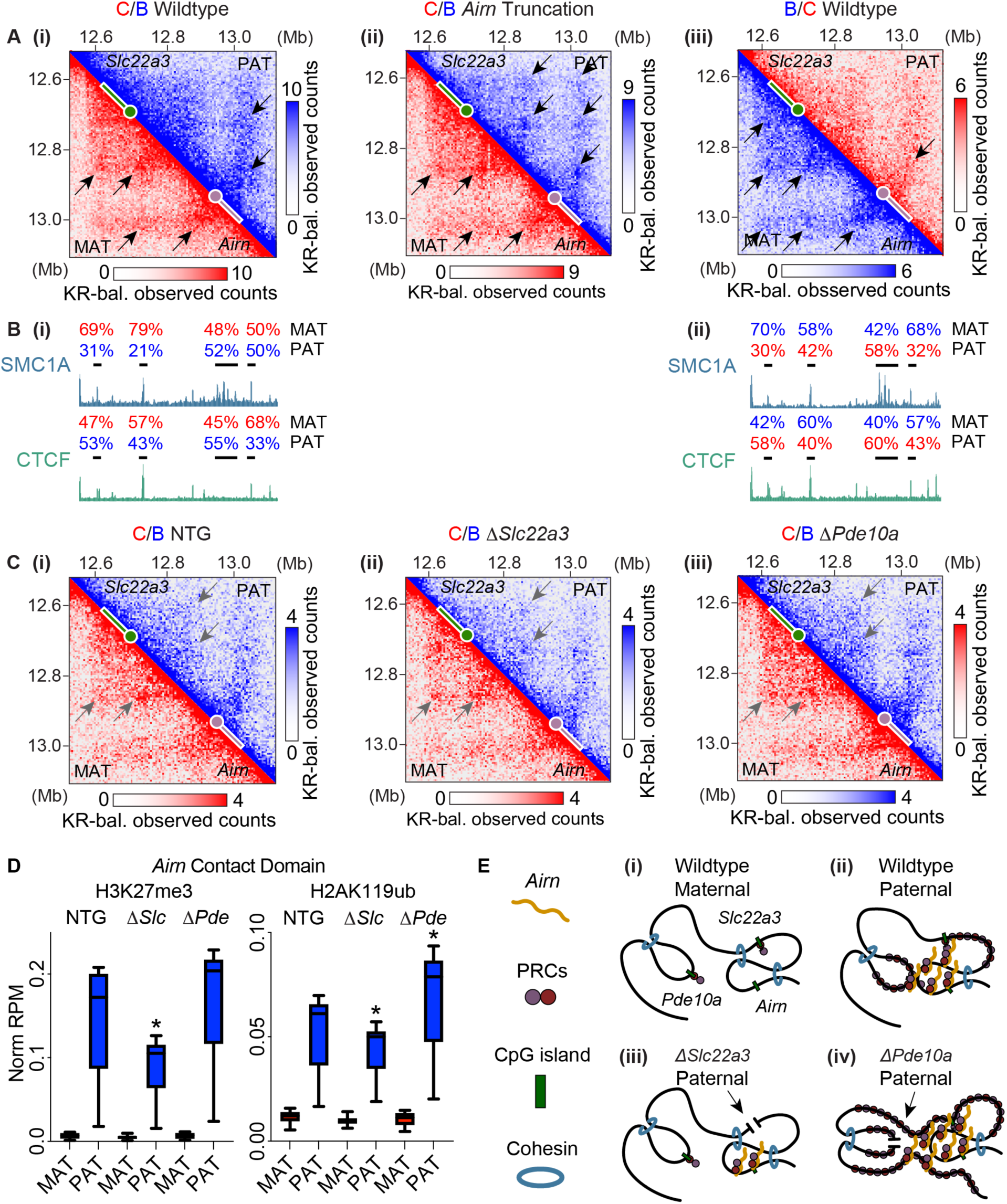
***Airn* expression is coincident with dissolution of DNA loops encasing *Slc22a3* and a local increase in PRC-directed modifications.** (A) Hi-C contact heatmaps of the *Airn* contact domain at 5kb resolution in (i) C/B wildtype, (ii) C/B *Airn* truncation, and (iii) B/C wildtype TSCs. Data processed as in Figure 1. KR-bal., Knight-Ruiz balanced. Black arrows, location of DNA loops detected by SIP. (B) ChIP-Seq density tracks for SMC1A/Cohesin and CTCF annotated with percent parental biases at loops, (i and ii) as in (A). (C) Contact heatmaps of the *Airn* contact domain at 5kb resolution in (i) C/B NTG, (ii) C/B Δ*Slc22a3*, and (iii) C/B Δ*Pde10a* TSCs. Data processed as in Figure 1. Grey arrows, location of DNA loops of interest from (A). In all heatmaps: circles, 5’ end of genes and CGIs; rectangles, gene bodies, purple, *Airn* gene; green, *Slc22a3* gene. (D) Boxplots of allelic H3K27me3 (left) and H2AK119 (right) ChIP-Seq density over the *Airn* contact domain in C/B NTG, C/B Δ*Slc22a3*, and C/B Δ*Pde10a* TSCs. Asterisks, significant changes relative to NTG (p<0.05, Welch t-test). (E) Model: DNA regulatory elements modulate frequency of contact with and repression by *Airn*. (i) On the maternal allele where *Airn* is not expressed, pre-existing contacts with *Airn* render certain regions more susceptible to repression than others. (ii) On the *Airn*-expressing paternal allele, PRCs and their modifications to chromatin potentiate contacts with *Airn*, which potentiates repression. (iii) Loss of *Slc22a3* CGI attenuates repression by reducing local recruitment of PRCs and contact with distal regions. (iv) Loss of *Pde10a* CGI increases the frequency with which surrounding regions contact *Airn*.

To investigate this possibility, we searched the NTG, Δ*Slc22a3*, and Δ*Pde10a* Hi-C data for DNA loops in the *Airn* contact domain. While lower sequencing depth (∼600 million read pairs per genotype) precluded a high-confidence analysis, visual inspection of the SIP-identified loops from our high-depth datasets in Figure 7A was consistent with the notion that the *Slc22a3* CGI is required for *Airn* to dissolve the nested loops encasing *Slc22a3* (Figure 7C; note relative increase in intensity at loop anchor regions in the *ΔSlc22a3* genotype, denoted by the two grey arrows). Also, we observed that deletion of the *Slc22a3* CGI led to a striking local drop in H3K27me3 and had a lesser yet significant effect on H2AK119ub (Figure 7D). Thus, in contrast to the *Pde10a* CGI, which may effectively restrict surrounding regions from contacting *Airn* through its own high- frequency contacts with the *Airn* locus, the *Slc22a3* CGI may enable *Airn*-recruited PRCs to engage with nearby chromatin more effectively, thereby displacing Cohesin, disrupting loops, and enabling *Airn* to contact distal regions more readily.

## DISCUSSION

Through a convergent analysis of Hi-C, ChIP-Seq, CHART-Seq, and RNA-Seq in wildtype and genetically perturbed F1-hybrid mouse TSCs and ESCs, we uncovered a series of striking relationships between 3D DNA contacts, DNA regulatory elements, and PRC recruitment within the largest autosomal region known to be repressed by a mammalian lncRNA. Our results strongly support the view that *Airn* is a potent *cis*-acting lncRNA whose predominant functions are to repress gene expression and recruit the PRCs to modify chromatin within a 15Mb genomic domain. Moreover, we show that the extent of repression induced by *Airn* can be modulated by discrete DNA regulatory elements that control the proximity of *Airn* to its genomic targets, a paradigm that we believe is relevant to other domains governed by strong locus control regions, including the inactive X chromosome and domains that surround super-enhancers (Hnisz et al. 2013; Arnold et al. 2019; Statello et al. 2021).

Using *in situ* Hi-C, we observed that expression of *Airn* is accompanied by dramatic changes in 3D contacts and compartmentalization on the centromeric side of the *Airn* locus. These changes correlated in-step with the intensity of *Airn*-induced H3K27me3 and H2AK119ub, and centered around three regions that contact the *Airn* locus even in the absence of *Airn* expression. Each of these regions harbor CGI promoters that bind components of vPRC1, exhibit signatures of transcriptional activity, and are located proximal to peaks of Cohesin on both the maternal and paternal alleles. Frequency of contact with the *Airn* lncRNA, as assessed by CHART-Seq, also correlated in-step with the intensity of PRC-directed modifications and centered around pre-existing contacts, in both TSCs and ESCs. Two genes – *Map3k4* and *Qk* – resisted accumulating PRC-directed modifications, despite both genes being repressed by *Airn* and forming contacts with the *Airn* locus. Moreover, the intensity of DNA contacts between *Map3k4*, *Qk*, and the *Airn* locus were relatively unchanged by *Airn* expression, and relative to surrounding regions, *Map3k4* and *Qk* resisted associations with the *Airn* lncRNA.

Together, our data support the notion that spatial proximity to the *Airn* lncRNA product induces gene repression, and in most cases, the accumulation of PRC-directed chromatin modifications. *Airn* is a short-lived RNA that does not diffuse away from its site of transcription (Seidl et al. 2006; Schertzer et al. 2019a). Thus, it follows that the regions that are most sensitive to repression by *Airn* are predisposed to contacting the *Airn* locus even in the absence of *Airn* expression. In turn, considering our own findings along with prior data showing that the PRCs and their modifications to chromatin can induce DNA compaction (Terranova et al. 2008; Rao et al. 2014; Blackledge and Klose 2021), it seems likely that *Airn*-recruited PRCs and the modifications they deposit on chromatin are responsible for the major changes in chromatin architecture induced by *Airn* expression. Such changes would presumably potentiate *Airn*-dependent silencing, stabilizing the process by positive feedback. *Airn*-induced repression of at least two genes, *Map3k4* and *Qk*, was observed in the absence of accumulated PRC-deposited modifications and associated changes in chromatin architecture. These data invoke the possibility that like *Xist*, *Airn* may initially repress genes using a pathway upstream of the PRCs, which requires the protein SPEN (Zylicz et al. 2019; Dossin et al. 2020). Those same data support the notion that PRC-directed modifications are responsible for the major changes in chromatin architecture observed upon expression of *Airn*.

Three notable regions within the *Airn* target domain – encompassing the genes *Slc22a3*, *Qk*, *Prr18/T/Pde10a* – exhibited augmented contacts with the *Airn* locus even in the absence of *Airn* expression. Of these regions, only *Slc22a3* formed a detectable DNA loop anchored at the *Airn* locus (Rowley et al. 2020). However, all of the regions harbored CGI promoters which themselves were associated with vPRC1, signatures of transcriptional activity, and nearby peaks of Cohesin.

Prior works would suggest that any or all of these features could facilitate interactions between the regions and *Airn* DNA (Rao et al. 2014; Boyle et al. 2020; Mirny and Dekker 2022). Upon expression of *Airn*, the region encompassing *Prr18/T/Pde10a* exhibited peak-like increases in contact with the *Airn* locus, while the intensity of detectable contacts with *Qk* and *Slc22a3* remained relatively unchanged. Deletion of specific DNA regulatory elements within the *Prr18/T/Pde10a* and *Slc22a3* regions affected the ability of *Airn* to induce gene silencing, PRC-directed modifications, and changes to chromatin architecture over multiple megabases. Thus, DNA elements shape long-range contacts within the *Airn* target domain in ways that extend beyond single loop-based models of regulation (Mirny and Dekker 2022).

Specifically, while deletion of the CGI at *T* had little to no effect, deletion of a ∼2.5kb region encompassing the *Pde10a* CGI or a ∼190kb cluster of CTCF/Cohesin peaks upstream of *Pde10a* both resulted in a striking increase in repression by *Airn* over the entirety of its target domain. Hi-C in Δ*Pde10a* cells showed a decreased interaction between *Airn* and *Pde10a* concomitant with an increased interaction between *Airn* and the remainder of its target domain. Moreover, relative to controls, the overall expression of the *Pde10a* gene appeared reduced in Δ*Pde10a* and Δ*Cluster* TSCs (Table S2). We speculate that high-frequency interaction between the *Airn* locus and the *Pde10a* CGI may dampen *Airn*’s ability to interact with surrounding DNA, and that disrupting this interaction by deletion enables *Airn* to interact more readily with other loci. The act of *Pde10a* transcription may also attenuate *Airn*’s ability to interact with surrounding DNA in a way that prevents widespread accumulation of PRC-directed modifications, even over distal regions. In either case, our data provide a remarkable example of how loss of a single DNA regulatory element can affect gene expression, chromatin modifications, and chromatin architecture over a region that is 15Mb in length, 7500 times longer than the deleted element itself.

Likewise, in a previous study, we had observed that deletion of the *Slc22a3* CGI led to a precipitous drop in PRC-deposited modifications in the 4.5Mb region between *Prr18*/*T*/*Pde10a* and *Airn* (Schertzer et al. 2019a). Here, we offer a mechanistic explanation for this otherwise perplexing observation. By Hi-C, we found that *Slc22a3* and *Airn* reside in the same contact domain structure, and prior to *Airn* expression, are entwined in nested DNA loops anchored by CTCF and Cohesin. Upon *Airn* expression, Cohesin binding and loops around *Slc22a3* are reduced. When the *Slc22a3* CGI is deleted, resulting Hi-C data are consistent with the reformation of local DNA loops. Thus, in contrast to the *Pde10a* CGI, which may promote contacts with *Airn* directly, the *Slc22a3* CGI may do so indirectly, by promoting local accumulation of PRC-deposited modifications upon exposure to *Airn*. In turn, these modifications could disrupt nearby DNA loops (Cuadrado et al. 2019; Rhodes et al. 2020; Kriz et al. 2021) and enable *Airn* to contact distal chromatin more efficiently.

Considered together, our data indicate that the extent of repression across the *Airn* target domain is governed by an equilibratory network of DNA regulatory elements that through direct or indirect means, control spatial proximity to the *Airn* lncRNA product (Figure 7E). Shifting the equilibrium in either direction has consequences on gene expression, chromatin modifications, and chromatin architecture. Indeed, we identified one CGI that appears to promote certain long-range contacts while restricting others (*Pde10a*), and another CGI that promotes long-range contacts presumably by serving as a local Polycomb Response Element (*Slc22a3*; Figure 7E; (Hojfeldt et al. 2018)). Meanwhile, the CGI promoters of *Map3k4* and *Qk* give the appearance of serving as boundary elements that attenuate the local spread of silencing by *Airn*. Thus, variation in the genetic or epigenetic content of a DNA regulatory element has the potential to control gene expression by altering spatial equilibria between genes and locus control regions, be those repressors or enhancers (Hnisz et al. 2013; Arnold et al. 2019; Greenwald et al. 2019; Statello et al. 2021; Galupa et al. 2022; Zuin et al. 2022). In turn, unrecognized alterations to spatial equilibria that modulate contact with locus control regions may contribute to the challenge of assigning target genes to disease-associated SNPs (Gazal et al. 2022).

Lastly, repression of the *Airn*-target gene *Igf2r* is due to the act of *Airn* transcription and does not depend on the *Airn* lncRNA product (Latos et al. 2012). However, the mechanisms responsible for long-range repression by *Airn* remain unclear. Our observation that *Airn* CHART signal and PRC-directed chromatin modifications correlate in lock-step over a 15Mb domain in both TSCs and ESCs, together with data from (Nagano et al. 2008; Andergassen et al. 2019; Schertzer et al. 2019a), strongly support the notion that the *Airn* locus produces a potent *cis*-acting repressive lncRNA that recruits the PRCs and possibly other repressive enzymes to broad regions of chromatin, likely using mechanisms related to those used by *Xist* (Markaki et al. 2021). Models to explain these data that exclude a function for the *Airn* lncRNA product would now be at odds with the principles of parsimony. By extension, it is conceivable that like *Airn*, other chromatin-associated RNAs harbor an ability to repress transcription and recruit the PRCs, both locally and over long distances. It has long been noted that *Airn* is poorly spliced, exceptionally long, and interspersed with common repeats (Lyle et al. 2000; Seidl et al. 2006). We would hypothesize that these same features are hallmarks of other *cis-*acting repressive RNAs.

## ACKNOWLEDGEMENTS

This work was supported by NIH grant R01GM136819 (J.M.C.), UM1HG009375 (E.L.), and R35GM128645 (D.H.P.). A.K.B was supported by NIGMS training grant T32GM119999 and by NICHD award F31HD103370. J.B.T. was supported by T32CA217824. E.S.D. was supported by NIH training grant T32GM067553. J.M.D. was supported by R35GM124764.

## AUTHOR CONTRIBUTIONS

A.K.B. and J.M.C. conceived the study; A.K.B., M.D.S., A.O., J.B.T., and J.M.C. performed the experiments; A.K.B., M.D.S., E.S.D., and J.M.C. performed the analyses; J.M.D, D.H.P., and E.L. provided reagents and advice; and A.K.B. and J.M.C. wrote the paper with feedback from co-authors.

## Supplemental Figures

**Figure S1.**
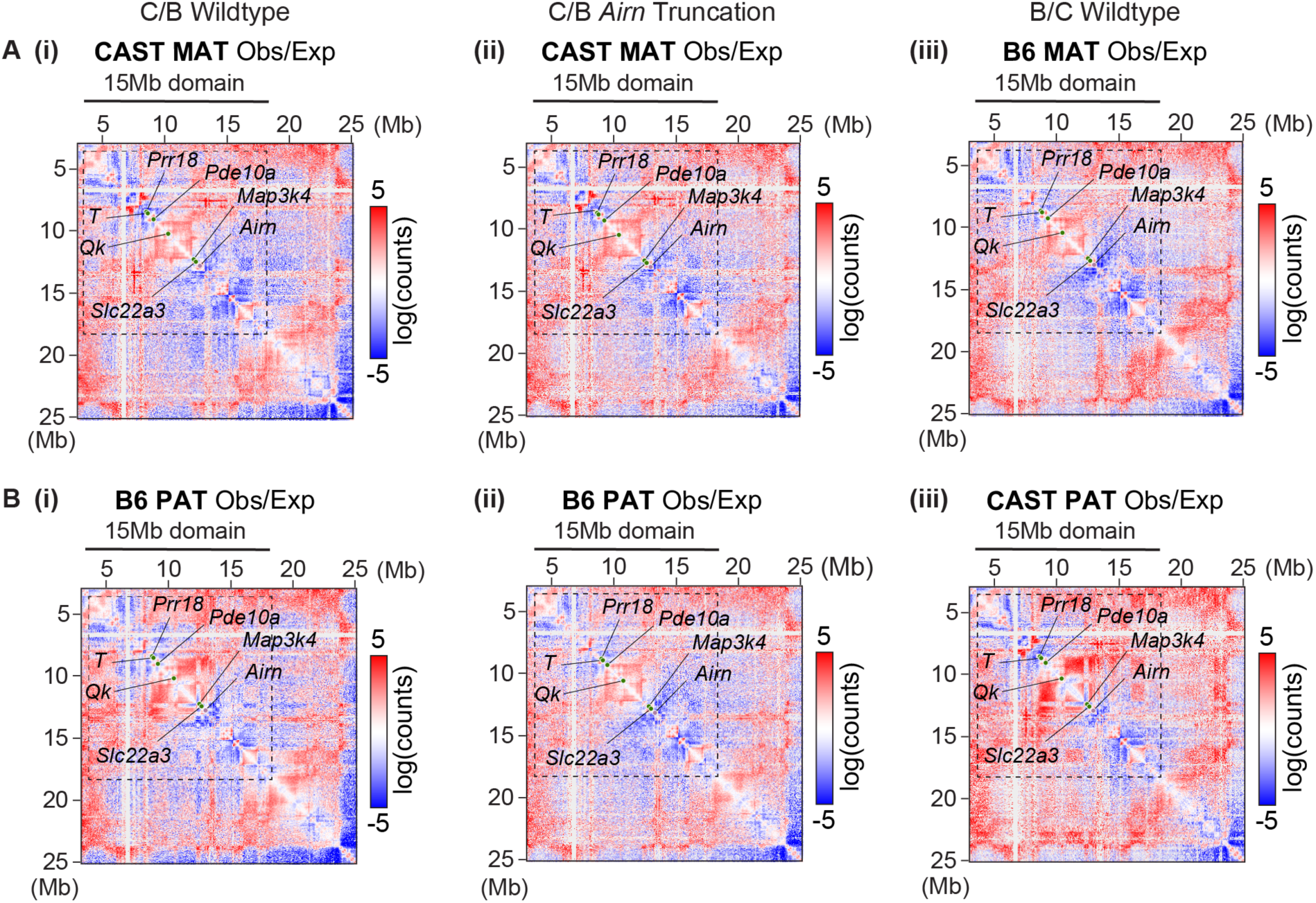
**(A, B)** 2D Hi-C contact heatmaps of maternal **(A)** and paternal **(B)** Observed-over- Expected (O/E) counts in **(i)** C/B wildtype, **(ii)** C/B *Airn* truncation, and **(iii)** B/C wildtype TSCs. Dotted lines, 15Mb *Airn* target domain; purple circle, *Airn* gene; green circles, other loci of interest.

**Figure S2.**
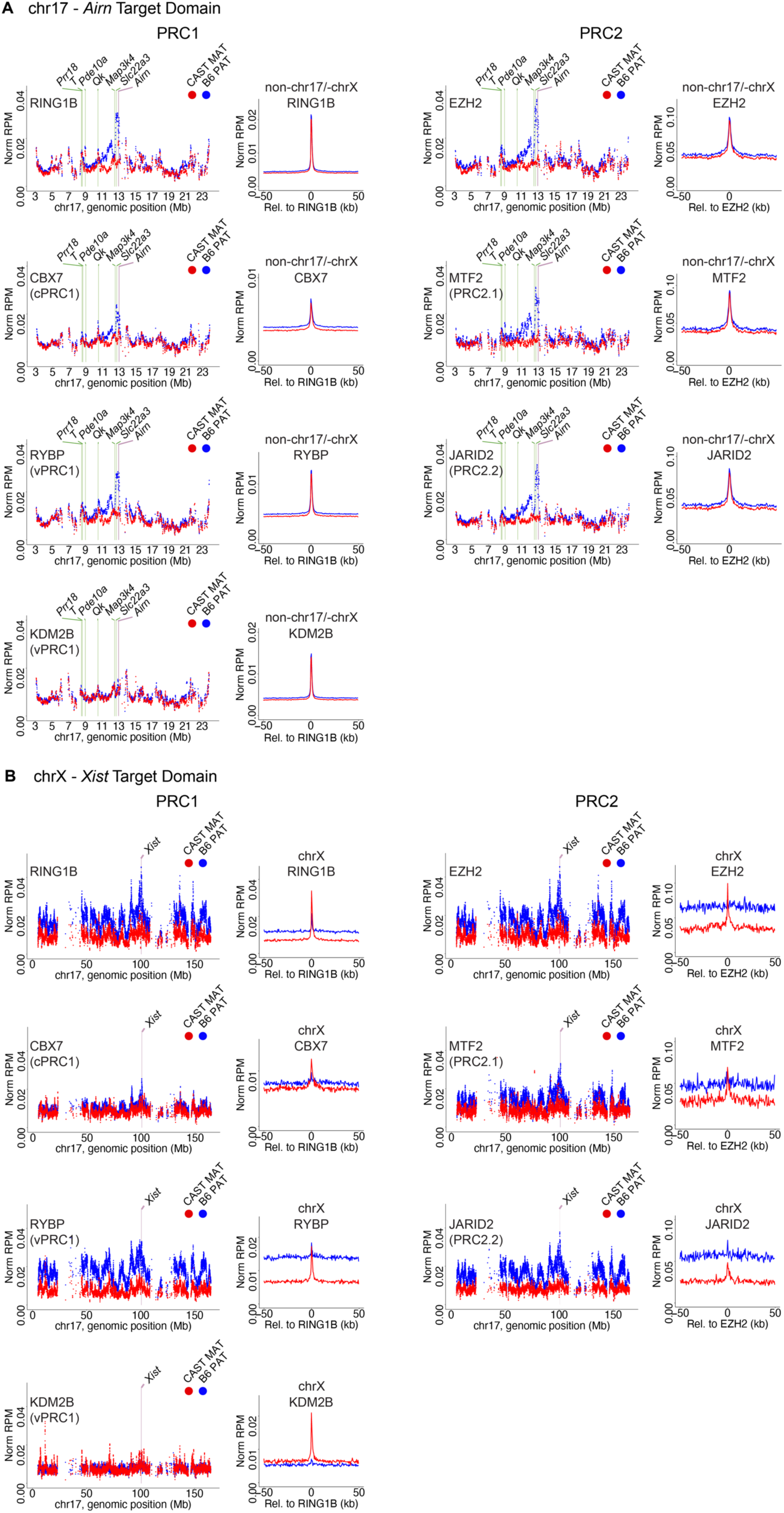
**(A)** Tiling density plots of allelic ChIP-Seq data for (left) PRC1 and (right) PRC2 components over the *Airn* target domain on chr17 in C/B TSCs. Data processed as in Figure 2. Shown to right, metagenes of SNP-normalized RPM values (Reads per Million total reads) for PRC subunits relative to all non-X, non-chr17 RING1B or EZH2 peaks. **(B)** Tiling density plots of allelic ChIP-Seq data for (left) PRC1 and (right) PRC2 components over chrX in C/B TSCs. Shown to right, metagenes of SNP-normalized RPM values for PRC subunits relative to RING1B or EZH2 peaks on chrX. In all panels: purple bar, *Airn* gene or *Xist* gene; green bars, other loci of interest.

**Figure S3.**
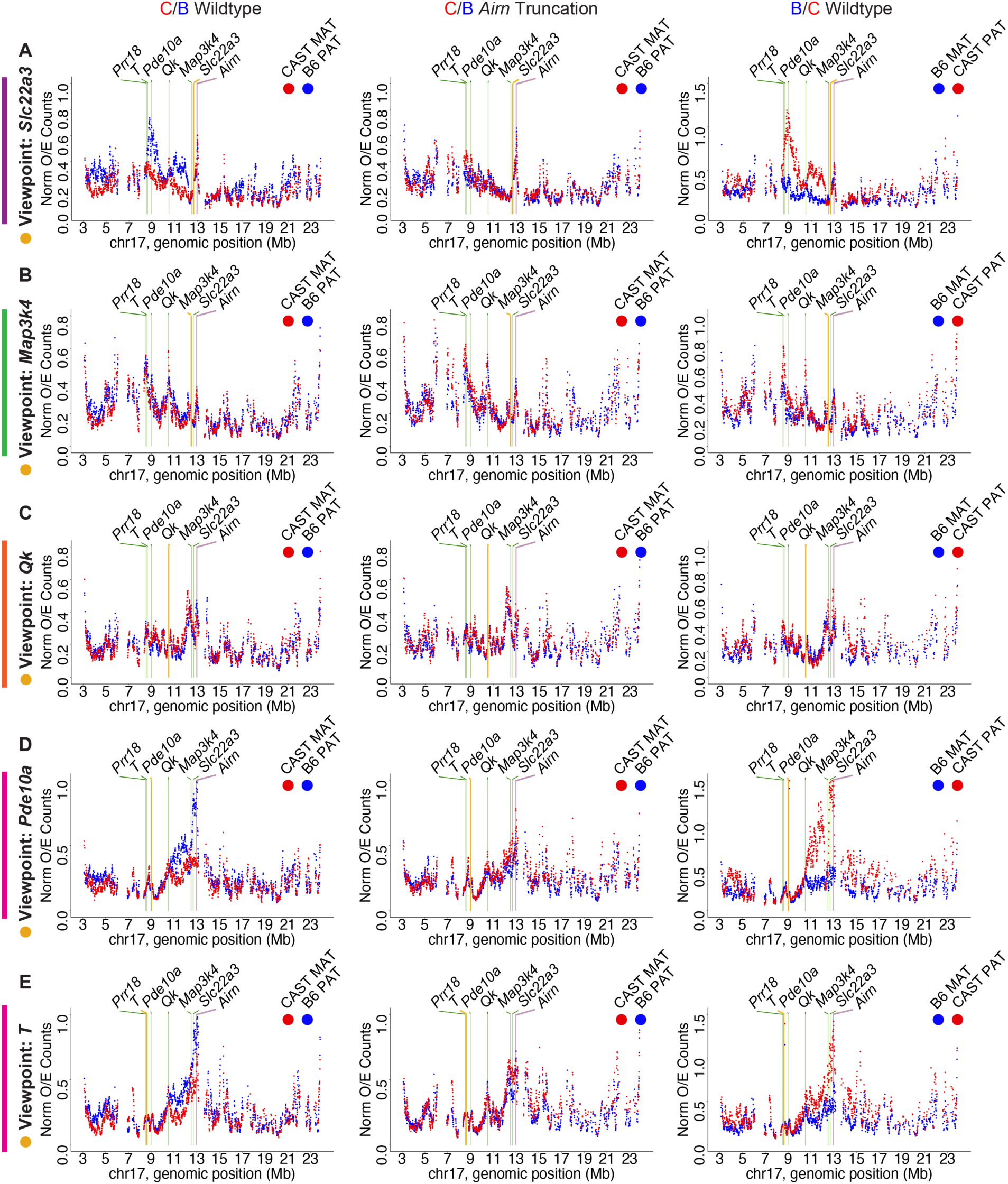
**(A-E)** Tiling density plots of allelic Hi-C Observed-over-Expected (O/E) counts for viewpoints of **(A)** *Slc22a3*, **(B)** *Map3k4*, **(C)** *Qk*, **(D)** *Pde10a*, and **(E)** *T* in **(i)** C/B wildtype, **(ii)** C/B *Airn* truncation, **(iii)** B/C wildtype TSCs. Data processed as in Figure 2. In all panels: yellow bar, viewpoint; purple bar, *Airn* gene; green bars, other loci of interest. Colored bars, regions analyzed in Figure 4.

**Figure S4.**
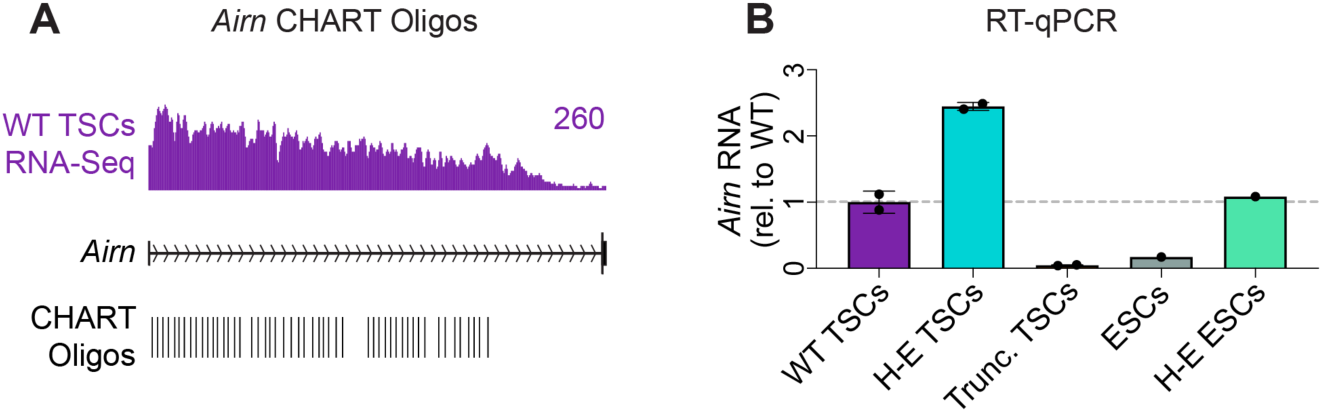
**(A)** Location of probes used for *Airn* CHART relative to an RNA-Seq density track of *Airn* in C/B wildtype TSCs. **(B)** RT-qPCR of *Airn* expression levels normalized by *Gapdh* and relative to WT TSCs.

**Figure S5.**
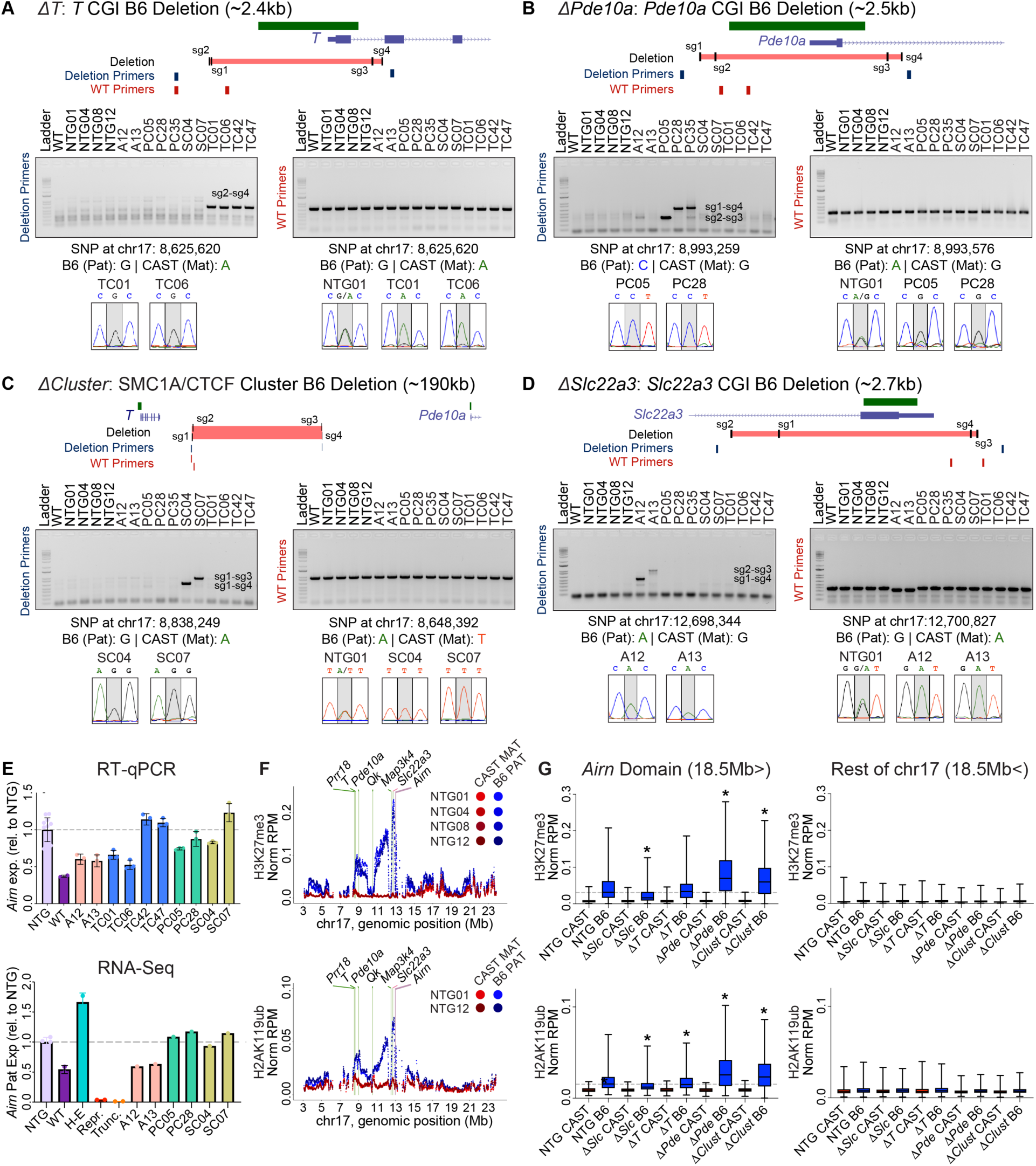
**(A-D)** Genotyping PCR gels and sequencing tracks for **(A)** Δ*T*, **(B)** Δ*Pde10a*, **(C)** Δ*Cluster*, and **(D)** Δ*Slc22a3* clonal TSC lines. Δ*Slc22a3* TSCs were generated in (Schertzer et al. 2019a). Gels of PCR product from either wildtype primers, which amplify the flanking end or internal region of the deletion site, or deletion primers, which amplify a sizeable region if the deletion occurred (Table S5). Sanger sequencing tracks of the PCR products show regions with B6/CAST SNPs for allelic identification. **(E)** RT-qPCR and RNA-Seq of *Airn* expression in clones relative to the averaged NTG value. **(F)** Tiling density plots of allelic H3K27me3 (top) and (bottom) H2AK119ub (bottom) ChIP-Seq data in C/B NTG TSCs. Data processed as in Figure 2. Purple bar, *Airn* gene; green bars, other loci of interest. **(E)** Boxplots of allelic H3K27me3 (top) and H2AK119 (bottom) ChIP-Seq density over the 15Mb *Airn* target domain (left) versus the rest of chr17 (right) in C/B NTG and deletion TSCs. Asterisks, statistically significant changes relative to NTG (p<0.05, Welch t-test).

**Figure S6.**
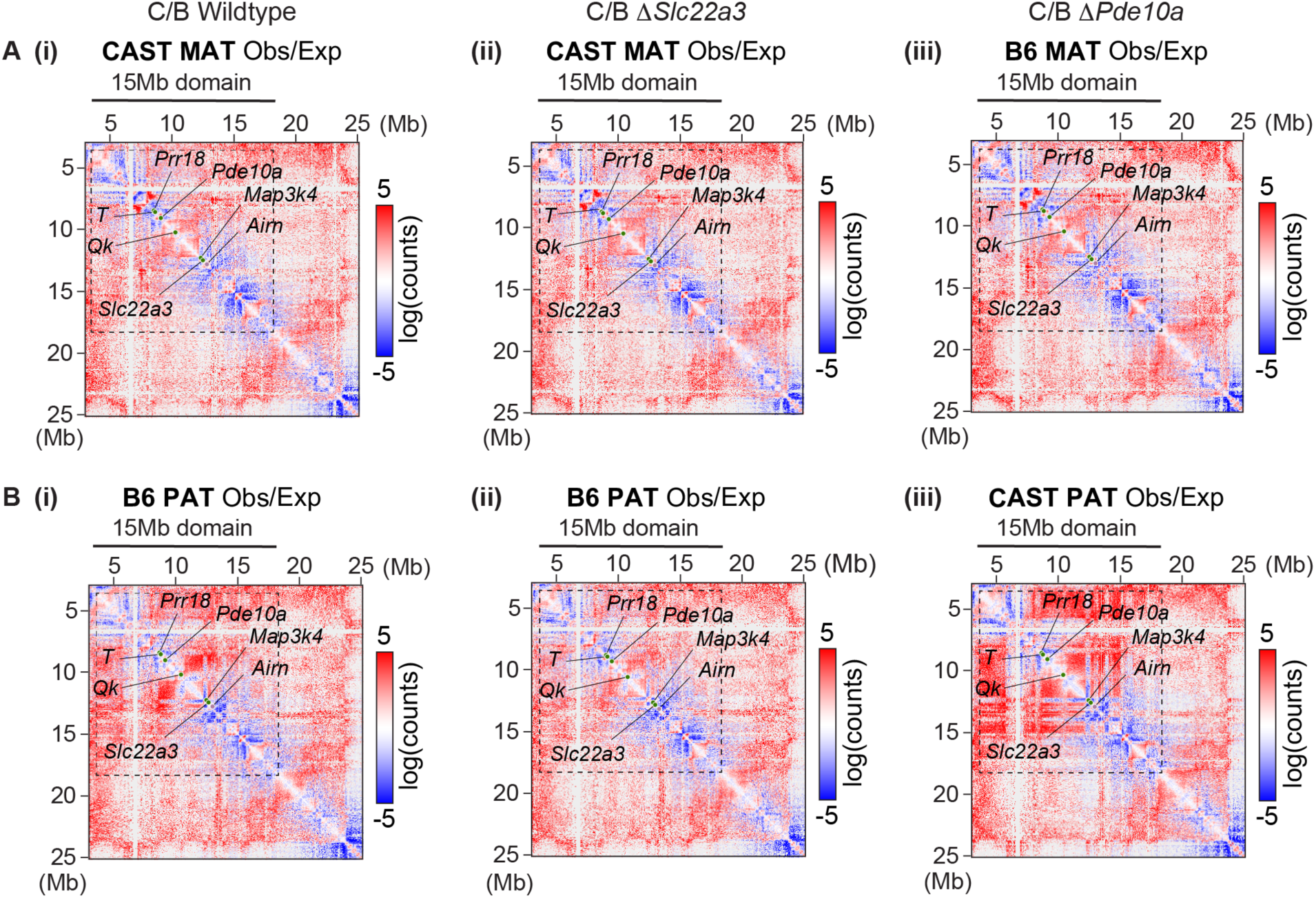
2D Hi-C contact heatmaps of maternal **(A)** and paternal **(B)** Observed-over-Expected (O/E) counts in **(i)** C/B NTG, **(ii)** C/B Δ*Slc22a3*, and **(iii)** C/B Δ*Pde10a* TSCs. Dotted lines, 15Mb *Airn* target domain; purple circle, *Airn* gene; green circles, other loci of interest.

## Supplemental Table Legends

**Table S1. Hi-C statistics and quality control.** Related to Figures 1, 2, 6, 7, S1, S3, S6. Table shows statistical summary of Hi-C datasets by total read depth, Juicer quality control statistics of Hi-C contacts, and HiCExplorer Pearson correlation of long-range contacts. Genotypes are separated by Sheets. Total read depth and Hi-C quality control sections were obtained from the Juicer output inter.txt (i.e., all read pairs) and inter_30.txt (i.e., MAPQ≥30 read pairs) files and referenced to the standard guidelines in (Rao et al. 2014). Pearson correlations of long-range Hi-C contacts (>25kb) at 10 and 25kb resolutions were determined by Hi-C Explorer’s hicCorrelate tool (Wolff et al. 2018).

**Table S2. RNA-Seq gene expression changes of *Airn* target genes.** Related to Figures 5E and S5E. Table shows gene expression changes across genotypes measured via RNA-Seq. Gene categories, separated by Sheets, include all chromosomes (‘All Chr’), the 27 genes in the *Airn* target domain that significantly change between *Airn* highly-expressing and *Airn* truncation TSCs (‘Airn Target Genes’; from (Schertzer et al. 2019a)), and the CGI-promoter genes of interest (‘CGI Genes’). All gene annotations are from gencode.vM1.annotation.gtf (Frankish et al. 2021). Reads aligning to introns are included. For total, non-allelic expression analysis, featureCounts (Liao et al. 2014) was used to count reads over exon coordinates (i.e., Counts, Total), and then divided by total reads in the dataset and multiplied by a million (i.e., RPM, Total). For allelic expression analysis, a custom perl script (ase_analyzer10.pl; see github) was used to count B6- and CAST SNP-overlapping reads over gene coordinates (i.e., Counts, CAST and B6).

**Table S3. Allelic detection of PRC components, architectural factors, and epigenetic marks.** Related to Figure 3B. Table shows the statistical summary of a permutation test used to determine whether factors of interest were significantly enriched at loci of interest relative to dataset specific background. All ChIP-Seq data were acquired from C/B TSCs. H3K27ac, H3K4me2, CTCF, and SMC1A data were generated in previous studies (Calabrese et al. 2012; Schertzer et al. 2019a). If applicable, all genomic features of interest were standardized to 1.5kb lengths (i.e., the largest CGI of interest) relative to their center positions. A list of 80,000 1.5kb regions were randomly selected from within ‘gene’ coordinates from gencode.vM1.annotation.gtf (Frankish et al. 2021) with 100kb extended start and end sites while excluding any regions that fell within 2.5kb of a region annotated by MACS as an H3K27me3 or PRC subunit peak. Shuffled coordinates were then filtered to retain regions encompassing at least one B6/CAST SNP, leaving 67,262 shuffled regions. B6- and CAST-overlapping ChIP-Seq reads were then counted over the features of interest and shuffled coordinates using a custom script (ase_analyzer10_adj2.pl; see github), then divided by the number of B6/CAST SNPs detected in the genomic coordinates (SNP-norm counts). The features were then ranked by SNP-norm counts for each allele in each dataset (1 = highest allelic signal), and a percentile rank was used to determine an empirical p-value for allele-specific enrichment of the ChIP target at the loci of interest. The raw allelic counts, rank, and empirical p-values for the genomic features of interest relative to the shuffled regions are separated by Sheets.

**Table S4. All high throughput sequencing datasets used.** Related to Figures 1-7, S1-S5 and Tables S1, S2, S3. Table gives all high throughput sequencing genomic datasets used in this study and is divided into 2 sections: “Datasets generated in this study” and “Publicly available datasets”. Under each section, if applicable: “File ID” gives the name of the dataset; “Experiment” gives laboratory experiment references; “Sequencing Date” gives the date the samples were loaded onto the sequencer, “Data Type” gives the experimental method (RNA-Seq, ChIP-Seq, CHART-Seq, Hi-C); “Cell Type” gives the cell type and strain information when relevant; “Spike-in” says whether ERCC or Drosophila spike-ins were included; “Read Length” describes 75bp versus 150bp read length and single versus paired end sequencing; “Figures and Tables” lists the figures and tables in the manuscript where each dataset was used; and “GEO” gives the GEO database reference for the data.

**Table S5. Oligonucleotides used.** Related to Figures 2, 4, 5, 6, 7, S4-6. Table gives all oligonucleotide sequences used in this study. “ID” gives a descriptive name for the oligonucleotide; “Assay” describes the experimental method in which the oligonucleotide was used (PCR genotyping, qPCR, CRISPR); “Sequence (5’->3’)” gives the oligonucleotide sequence in 5’ to 3’ order; and “Usage” gives the figure where the oligonucleotide was used.

## STAR Methods

### KEY RESOURCES TABLE

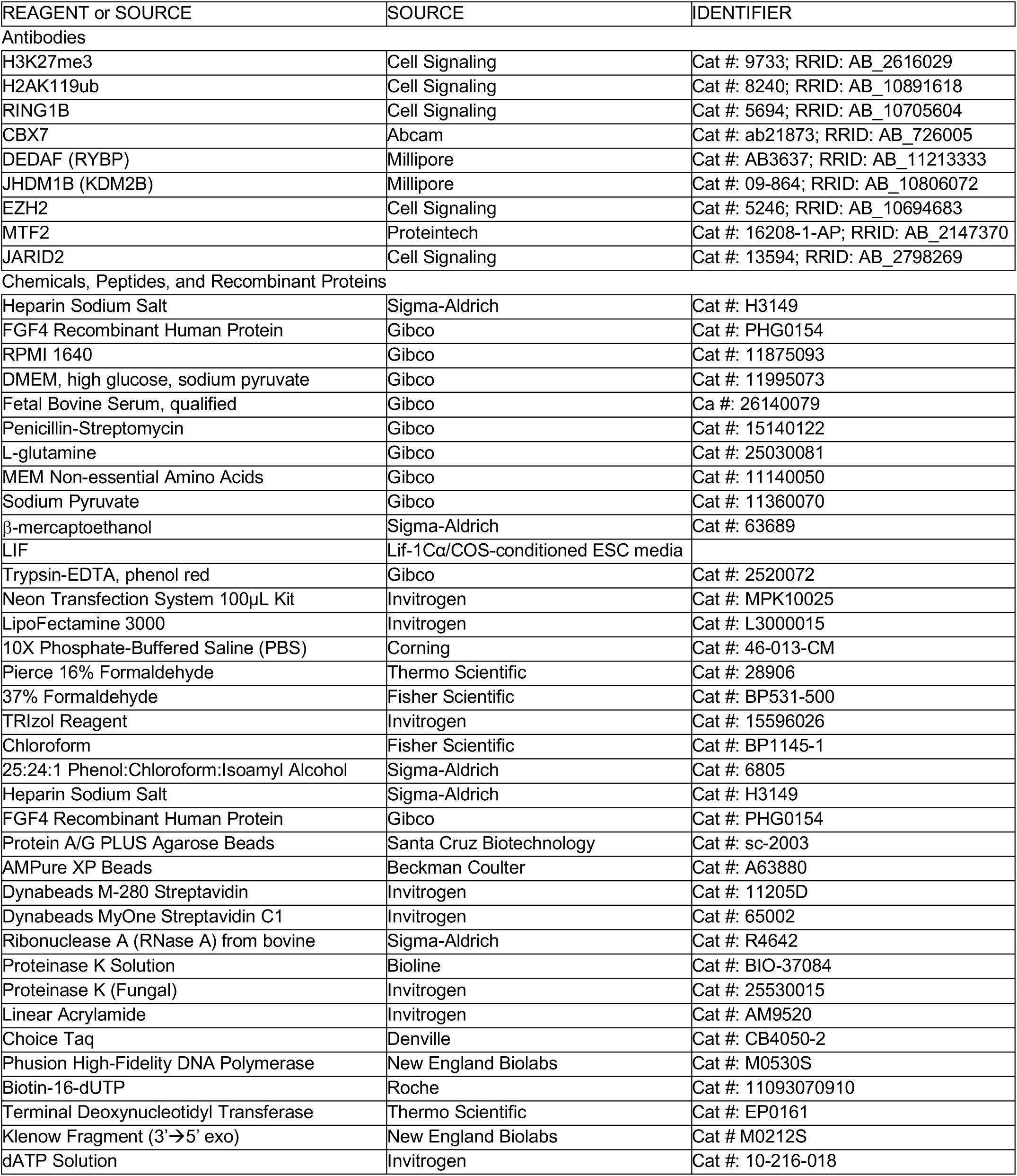
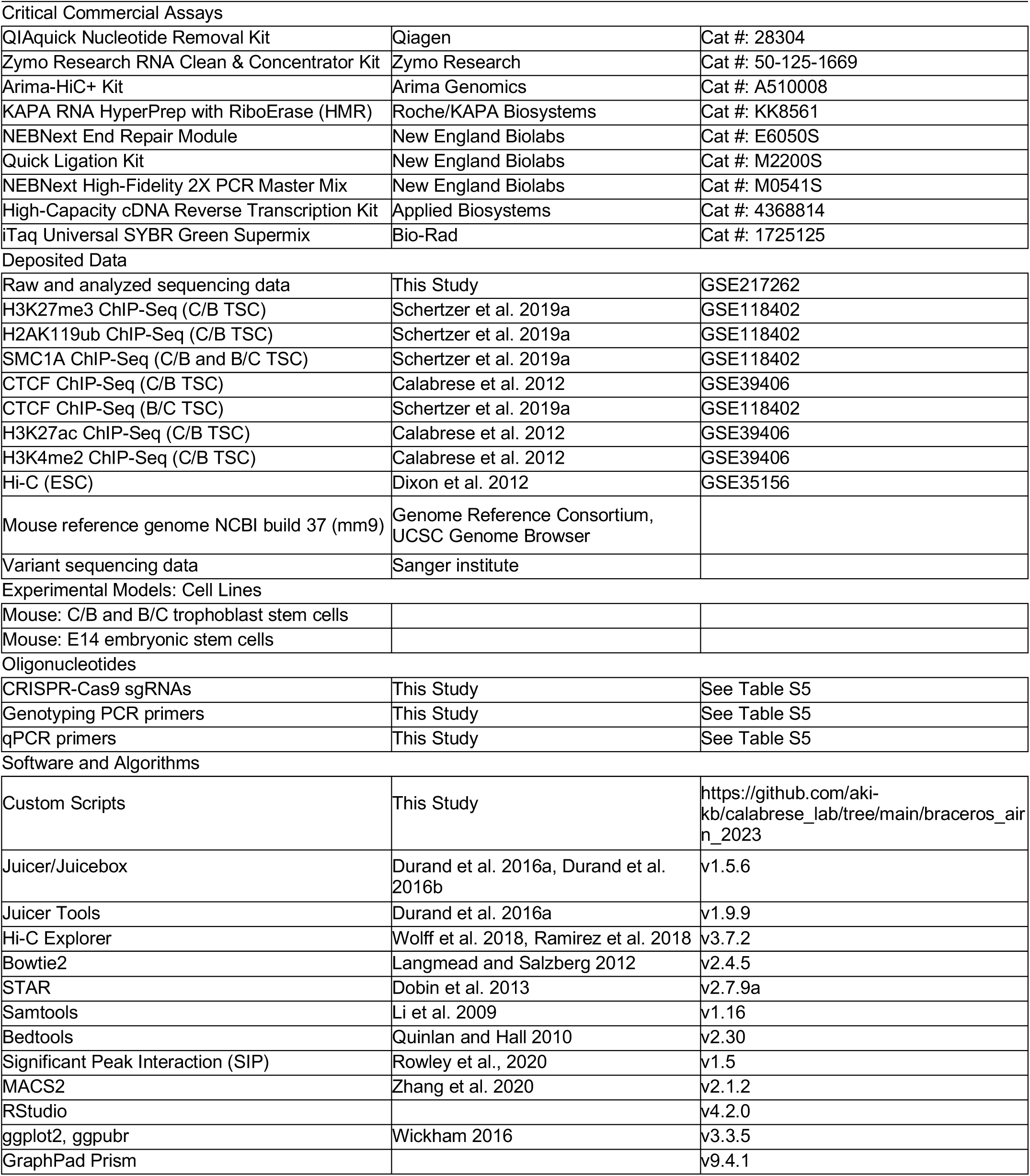

## CONTACT FOR REAGENT AND RESOURCE SHARING

Further information and requests for resources and reagents should be directed to and will be fulfilled by the Lead Contact, Mauro Calabrese.

## EXPERIMENTAL MODEL AND SUBJECT DETAILS

### Trophoblast stem cell (TSC) culture derivation and culture

The mouse C/B TSC and B/C TSC lines used in this work correspond to the CAST/EiJ maternal/C57BL/6J paternal (C/B) and C57BL/6J maternal/CAST/EiJ paternal (B/C) TSCs used in (Calabrese et al. 2012) and (Schertzer et al. 2019), and are referred to as CB.1 and BC.1 TSCs in (Calabrese et al. 2015). TSCs were cultured as in (Quinn et al. 2006). Briefly, TSCs were cultured on gelatin-coated, pre-plated irradiated mouse embryonic fibroblast (irMEF) feeder cells in TSC media (RPMI [Gibco, cat #: 11875093], 20% qualified FBS [Gibco, cat #: 26140079], 0.1mM penicillin-streptomycin [Gibco, cat #: 15140122], 1mM sodium pyruvate [Gibco, cat #: 11360070], 2mM L-glutamine [Gibco, cat #: 25030081], 100μM β-mercaptoethanol [Sigma-Aldrich, cat #: 63689]) supplemented with 25ng/mL FGF4 (Gibco, cat #: PHG0154) and 1μg/mL Heparin (Sigma-Aldrich, cat #: H3149) just before use, at 37°C in a humidified incubator at 5% CO2. At passage, TSCs were trypsinized with 0.125% Trypsin-EDTA in PBS solution (Gibco, cat #: 25200-072) for ∼4 minutes at room temperature and gently dislodged from the plate with a sterile, cotton-plugged Pasteur pipette. To deplete irMEFs from TSCs prior to all harvests, TSCs were pre-plated for 45 minutes at 37°C, transferred to a fresh culture plate, and then cultured for three days in 70% irMEF-conditioned TSC media supplemented with growth factors as above.

### Embryonic stem cell (ESC) culture

Mouse E14 ESCs were cultured on gelatin-coated plates in ESC media (DMEM high glucose and sodium pyruvate [Gibco, cat #: 11995073], 15% qualified FBS, 0.1mM MEM non-essential AA [Gibco, cat #: 11140050], 0.1mM penicillin-streptomycin, 2mM L-glutamine, 100μM β-mercaptoethanol, 1:500 LIF) at 37°C in a humidified incubator with 5% CO_2_. At passage, ESCs were trypsinized with 0.125% Trypsin-EDTA in PBS solution for ∼5 minutes at room temperature and dislodged from the plate at single-cell suspension. ESCs were passaged every other day and provided fresh media daily.

### Generation of clonal TSCs with DNA regulatory element deletions

Per regulatory element deletion, four unique sgRNAs were designed using CRISPOR (Concordet and Haeussler 2018), with two sgRNAs flanking the target site (Figure S2A-D, Table S5). As a negative control, an sgRNA using a non-targeting (NTG) sequence from (Invitrogen, cat #: A35526) was designed. Each sgRNA was cloned into the *BsmbI* site of the piggyBac-cargo rtTA vector from (Schertzer et al. 2019b) and transformed in DH5-alpha competent bacterial cells. Starter transformant cultures for each sequence-verified sgRNA were pooled together in equal volume amounts prior to liquid culture expansion and plasmid purification using the PureLink HiPure Plasmid Midiprep kit (Invitrogen, cat #: K2100004). The pooled sgRNAs were then co-electroporated with doxycycline-inducible Cas9-cargo and pUC19-piggyBac transposase vectors from (Schertzer et al. 2019b) at an 8:2:1 plasmid ratio of 2.5µg total DNA into 1 million C/B TSCs on irradiated drug-resistant MEFs (irDR4-MEFs; ATCC, cat #: SCRC-1045) in a single well of a 6-well plate. The electroporations were performed using a Neon® Instrument (electroporation settings: 950V, 30ms, 2 pulses). Two days after electroporation, TSCs were selected with 150µg/mL hygromycin B (Corning, cat #: MT30240CR)) and 200µg/mL G418 (Gibco, cat #: 10131035) in irMEF-conditioned TSC media with growth factors for 11 days, followed by four days of 1µg/mL doxycycline treatment (Sigma-Aldrich, cat #: D9891) to induce Cas9 expression. 2,000 doxycycline-induced TSCs were then plated onto a pre-plated irMEFs 100-mm dish for clonal selection and expansion. Prior to harvesting for genotyping assays, clonal TSC lines were passaged once off of irMEFs as above.

For genotyping assays, PCR was used to detect the presence or absence of target deletion DNA. “Wildtype” primers were designed to amplify either the flanking end or internal region of the deletion site. “Deletion” primers were designed to externally flank both ends of the deletion sites that would efficiently amplify a sizeable PCR product if the deletion occurred (Table S5). Sanger sequencing (Eton) was then used to detect the presence of informative B6/CAST SNPs in the PCR products for allelic identification.

### Generation of *Airn*-overexpressing ESCs

750,000 ESCs were seeded in a single gelatin-coated well of a six-well plate, and the next day transfected with 2.5µg of an 8:2:1 plasmid ratio of piggyBac-cargo rtTA-*Airn* sgRNA, doxycycline-inducible piggyBac-cargo dCas9-VP160, and pUC19-piggyBac transposase vectors from (Schertzer et al. 2019b) using Lipofectamine 3000 (Invitrogen, cat #: L3000015) according to manufacturer instructions. The next day, transfected ESCs were selected with ESC media containing 150µg/mL hygromycin B and 200µg/mL G418 for 10 days, followed by 4 days of doxycycline treatment to induce *Airn* expression via dCas9-VP160 prior to harvests.

## EXPERIMENTAL METHOD DETAILS

### In situ Hi-C

Prior to crosslinking for Hi-C, TSCs were passaged once off of irMEFs as described above. TSCs were then trypsinized and washed once with PBS. 5-10 million cells were crosslinked in resuspension with 10mL of 1% formaldehyde (Thermo Scientific, cat #: 28906) in PBS solution for 10 minutes at room temperature, quenched with 200mM glycine for 5 minutes at room temperature, and then washed twice with ice-cold PBS. Cells were then divided into 5 aliquots (1- 2 million cells/aliquot), where one aliquot was used for each Hi-C experiment. Importantly, for the removal of all PBS washes and crosslinking solution, cells were spun at 160 x g for 5 minutes.

Hi-C libraries from C/B wildtype, B/C wildtype, and C/B *Airn* truncation TSCs were generated and sequenced as in the detailed protocol from (Rao et al. 2014), including DNA fragmentation with *MboI* and *MseI* restriction enzymes. Hi-C libraries from regulatory element deletion TSCs were generated using the Arima-HiC+ kit (Arima Genomics, cat #: A510008) according to the manufacturer instructions. Paired-end, 150-bp sequencing was performed using Illumina NovaSeq 6000 System.

### ChIP-Seq

Prior to crosslinking for ChIP, TSCs were passaged once off of irMEFs as above. For all ChIP experiments, except those for PRC components, adhered cells were crosslinked with 0.6% formaldehyde (Fisher Scientific, cat #: BP531-500) in RPMI media with 10% FBS for 10 minutes at room temperature, then quenched with 125mM glycine for 5 minutes at room temperature. Crosslinked cells were then washed twice with ice-cold PBS and scraped with ice-cold PBS with 0.05% Tween (Fisher Scientific, cat #: EW-88065-31) and PIC (Sigma Aldrich, cat #: P8340). The cells were then spun at 1,200 x g at 4°C to remove PBS, followed by resuspension in ice-cold PBS with PIC and divided into 10-million cell aliquots. For PRC component ChIPs, adhered C/B TSCs were crosslinked in PBS with 2mM DSG (disuccinimidyl glutarate; Thermo Scientific, cat #: 20593) for 45 minutes at room temperature and then in PBS with 1% formaldehyde (Thermo Scientific, cat #: 28906) for 15 minutes at room temperature. Crosslinking was quenched with 200mM glycine for 5 minutes at room temperature. Cells were then washed, scraped, and aliquoted as above. All ChIPs were performed using 10 million cells, 10μL of antibody, and 30μL of Protein A/G agarose beads (Santa Cruz, cat #: sc-2003). Input chromatin was isolated accordingly to each antibody (see below) and sonicated to 100-500bp fragments using a Vibracell VX130 (Sonics) with the following parameters: 8-10 cycles of 30% intensity for 30 seconds with 1 minute of rest on ice between cycles. Antibody-conjugated beads were prepared by incubating antibody with beads in 300μL Blocking Buffer (PBS, 0.5% BSA [Invitrogen, cat #: AM2616]) overnight at 4°C with rotation.

For H3K27me3 and H2AK119ub ChIPs, crosslinked TSCs were resuspended in 1mL Lysis Buffer 1 (50mM HEPES pH 7.5, 140mM NaCl, 1mM EDTA, 10% glycerol, 0.5% NP-40, 0.25% Triton X-100, PIC) and incubated with rotation for 10 minutes at 4°C. Cells were then resuspended in 1mL Lysis Buffer 2 (10mM Tris-HCl pH 8.0, 200mM NaCl, 1mM EDTA, 0.5 mM EGTA, PIC) for 10 minutes at room temperature. All buffer removal steps were performed with 5- minute 1,200 x g spins at 4°C. The extracted nuclei pellet was then resuspended and sonicated in 500μL Lysis Buffer 3 (10mM Tris-HCl pH 8.0, 100mM NaCl, 1mM EDTA, 0.5mM EGTA, 0.1% sodium-deoxycholate, 0.5% N- lauroylsarcosine, PIC). Soluble chromatin was obtained with a 30- minute max speed spin at 4°C, mixed with 1% Triton X-100, and then incubated with pre- conjugated antibody beads overnight at 4°C with rotation. The ChIP beads were then washed five times in 1mL RIPA Buffer (50mM HEPES pH 7.5, 500mM LiCl, 1mM EDTA, 1% NP-40, 0.7% sodium-deoxycholate, PIC) and once with 1mL TE, each for 5 minutes at 4°C with rotation and spun at 2,000 x g for 2 minutes for buffer removal.

For PRC component ChIPs, crosslinked C/B TSCs were resuspended and sonicated in 500μL Low Salt Pol II ChIP Buffer (50mM Tris-HCl pH 7.5, 140mM NaCl, 1mM EDTA, 1mM EGTA, 0.1% sodium-deoxycholate, 0.1% SDS, PIC). Soluble chromatin was obtained with a 30-minute max speed spin at 4°C, mixed with 1% Triton X-100, and then incubated with pre-conjugated antibody beads overnight at 4°C with rotation. The ChIP beads were then washed three times with 1mL Low Salt Pol II ChIP Buffer with 1% Triton X-100 and PIC, once with 1mL High Salt Pol II ChIP Buffer (50mM Tris-HCl pH 7.5, 500mM NaCl, 1mM EDTA, 1mM EGTA, 0.1% sodium- deoxycholate, 0.1% SDS, PIC), once with 1mL LiCl Wash Buffer (20mM Tris-HCl pH 8.0, 250mM LiCl, 1mM EDTA, 0.5% Na-deoxycholate, 0.5% NP-40, PIC), and once with 1mL TE, each for 5 minutes at 4°C with rotation and spun at 2,000 x g for 2 minutes for buffer removal.

For RAD21 ChIP, crosslinked TSCs were resuspended and incubated with Lysis Buffer 1 and Lysis Buffer 2 as above. The extracted nuclei pellet was then resuspended and sonicated in 500μL Shearing Buffer (20mM Tris-HCl pH 8.0, 150mM NaCl, 2mM EDTA, 0.1% SDS, PIC). Soluble chromatin was obtained with a 30-minute max speed spin at 4°C, mixed with 1% Triton X-100 and 50mM NaCl, and then incubated with pre-conjugated antibody beads overnight at 4°C with rotation. ChIP beads were washed once with 1mL Lysis Buffer 3 (20mM Tris-HCl pH 8.0, 150 mM NaCl, 2mM EDTA pH 8.0, 0.1% SDS, 1% Triton X-100, PIC), once with 1mL High Salt Buffer C (20mM Tris-HCl pH 8.0, 500mM NaCl, 2mM EDTA pH 8.0, 0.1% SDS, 1% Triton X-100, PIC), once with 1mL Low Salt Buffer D (10mM Tris-HCl pH 8.0, 250 mM LiCl, 1mM EDTA pH 8.0, 1% NP-40, PIC), and once with 1mL TE with 50mM NaCl, each for 5 minutes at 4°C with rotation and spun at 2,000 x g for 2 minutes for buffer removal.

For all ChIP DNA elution steps, washed beads were resuspended in Elution buffer (50mM Tris-HCl pH 8.0, 10mM EDTA, 1% SDS) and placed on a 65°C heat block for 17 minutes with frequent vortexing. RAD21 ChIP DNA was eluted for 1 hour under the same conditions. ChIP DNA was then reverse crosslinked in 0.5% SDS and 100mM NaCl overnight at 65°C, followed by a 1-hour RNaseA (3μL; Thermo Scientific, cat #: EN0531) treatment at 37°C and a 2.5-hour Proteinase K (10μL; Invitrogen, cat #: 25530015) treatment at 56°C. DNA was then extracted with 1 volume of phenol:chloroform:isoamyl alcohol (Sigma-Aldrich, cat #: P3803) and precipitated with 2 volumes 100% ethanol, 1/10 volume 3M sodium-acetate pH 5.4, and 1/1000 volume linear acrylamide (Invitrogen, cat #: AM9520) overnight at -20°C. Precipitated DNA was then extracted with a 30-minute max speed spin at 4°C, washed once with ice-cold 80% ethanol, and resuspended in TE.

ChIP-Seq libraries were prepared with NEBNext End Repair Module (NEB, cat #: E6050S), A-tailing by Klenow Fragment (3’à5’ exo-; NEB, cat #: M0212S), and TruSeq 6-bp index adaptor ligation by Quick ligase (NEB, cat #: M2200S), and NEBNext High-Fidelity 2X PCR Master Mix (NEB, cat #: M0541S). All DNA clean-up steps were performed with AMPure XP beads (Beckman Coulter, cat #: A63880). Single-end, 75-bp sequencing was performed using an Illumina NextSeq 500/550 High Output v2.5 kit (Illumina, cat #: 20024906) on a NextSeq 500 System.

### CHART-Seq

CHART was performed as in the detailed protocol from (Davis and West 2015). TSCs were passaged once off of irMEFs as above. *Airn* Highly-Expressing TSCs from (Schertzer et al. 2019a) and ESCs were induced with 1μg/mL doxycycline 4 days prior to crosslinking. Adhered TSCs and ESCs were crosslinked with 1% formaldehyde (Fisher Scientific, cat #: BP531-500) in PBS solution for 10 minutes at room temperature. On ice, cells were washed twice with ice-cold PBS and twice with ice-cold PBS + 0.05% Tween before being scraped, spun at 1,000 x g for 5 minutes at 4°C, and divided into 25-million cell aliquots.

Per aliquot, nuclei was extracted with two rounds of douncing in 4mL sucrose buffer (300mM sucrose, 10mM HEPES pH 7.5, 100mM KOAc, 1% Triton X-100, 0.1mM EGTA, 0.5mM spermidine, 0.15mM spermine, cOmplete EDTA-free protease inhibitor cocktail [Millipore, cat #: 11873580001], 1mM DTT, 80U SUPERase-in [Invitrogen, cat #: AM2696]), mixing 1:1 with glycerol buffer (25% glycerol, 10mM HEPES pH 7.5, 100mM KOAc, 1mM EDTA, 0.1mM EGTA, 0.5mM spermidine, 0.15mM spermine, cOmplete EDTA-free protease inhibitor cocktail, 1mM DTT, 80U SUPERase-in), and centrifugation through 4mL glycerol buffer at 1,000 x g for 15 minutes at 4°C. The nuclei pellet was then washed twice with ice-cold PBS + 0.05% Tween, then further crosslinked with 3% formaldehyde (Fisher Scientific, cat #: BP531-500) in PBS + 0.05% Tween for 30 minutes at room temperature. Formaldehyde was then washed out twice with ice-cold PBS + 0.05% Tween, then resuspended in freshly prepared 250μL Sonication Buffer (50mM HEPES pH 7.5, 75mM NaCl, 0.5% *N*-lauroylsarcosine solution, 0.1% sodium-deoxycholate, 0.1mM EGTA, cOmplete EDTA-free protease inhibitor cocktail, 1mMDTT, 300U SUPERase-in). Chromatin was sonicated to 2-10kb fragments using a Bioruptor Plus sonication device (Diagenode; sonication parameters: 30 sec on, 30 sec off cycles on high setting), then spun at max speed for 30 minutes at 4°C to retrieve soluble chromatin.

For *Airn* CHART, we designed 22-nucleotide complementary oligos that tile across the first 75 kb of the *Airn* RNA sequence using the ChIRP Probe Designer (LGC Biosearch Technologies) under parameters of high masking for specificity and ≥500-nt spacings (Table S5). The resulting 51 oligo probes were then mixed to a 100μM pool for in-house oligo biotin labeling (Flickinger et al. 1992). Briefly, 20μM oligo probe mix was labeled with 300μM biotin-16-dUTP (Roche, cat #: 11093070910) and 30U terminal deoxynucleotidyl transferase (Thermo Scientific, cat #: EP0161), then labeled for 15 minutes at 37°C and inactivated for 20 minutes at 75°C. Biotinylated oligo probes were then purified using the QIAquick Nucleotide Removal kit (Qiagen, cat #: 28304), eluting to ∼20μM.

Per *Airn* CHART reaction, 12.5 million cells worth of chromatin was mixed with 0.5 volume PAB (8M Urea, 100mM HEPES pH 7.5, 200mM NaCl, 2% SDS) and 1.5 volumes of freshly prepared Hybridization Buffer (1.5M NaCl, 1.12M Urea, 10X Denhardt’s solution [Invitrogen, cat #: 750018], 10mM EDTA), then pre-cleared with 50μL Dynabeads M-280 Streptavidin beads (Invitrogen, cat #: 11205D) for 1 hour at room temperature with rotation. The pre-cleared chromatin was then isolated from the beads by spinning at 1,000 x g for 30 seconds. 1% of pre-cleared chromatin was saved as “Input” sample, and the remaining pre-cleared chromatin was incubated with 750pmol biotinylated oligo probes overnight at room temperature. The next day, the sample was spun at max speed for 20 minutes at 20°C, and the supernatant was incubated with 200μL worth of Dynabeads MyOne Streptavidin C1 beads (Invitrogen, cat #: 65001) resuspended in 125μL 2:1 diluted PAB overnight at room temperature with rotation. CHART beads were then placed on the magnet and washed 4 times with 400μL CHART Wash Buffer (250mM NaCl, 10mM HEPES pH 7.5, 2mM EDTA, 2mM EGTA, 0.2% SDS, 0.1% *N*-lauroylsarcosine) and once with RNase H Elution Buffer (75mM NaCl, 50mM HEPES pH 7.5, 0.02% sodium-deoxycholate, 0.1% *N*-lauroylsarcosine, 10mM DTT, 3mM MgCl2, 200U SUPERase-in). Beads were resuspended in 100μL RNase H Elution Buffer and treated with 2μL RNase H (NEB, cat #: M0297) for 10 mins at room temperature. To stop the RNase H reaction, 1/4 volume of naXLR (166.7mM Tris-HCl pH 7.2, 1.67% SDS, 83.3mM EDTA, 600μg Proteinase K [Bioline, cat #: BIO-37084]) was added, and the CHART eluate was then subject to proteinase K digestion and reverse-crosslinking for 1 hour at 55°C, followed by 1 hour at 65°C. The sample was then split for DNA (90%) and RNA (10%) analysis.

The *Airn* CHART-enriched RNA sample was treated with 1mL TRIZol and 200μL chloroform, and the RNA was DNase-treated and purified with Zymo Research RNA Clean & Concentrator kit (Zymo Research, cat #: 50-125-1669). To determine the extent of *Airn* RNA enrichment, 50% of both Input and CHART RNA samples were reverse transcribed using High-Capacity cDNA Reverse Transcription Kit (Applied Biosystems, cat #: 4368814). qPCR was performed using iTaq Universal SYBR Green Supermix (Bio-Rad, cat #: 1725125) and primers targeting 45 kb downstream of the *Airn* TSS and *Gapdh* (Table S5).

The *Airn* CHART-enriched DNA sample was extracted with 1 volume of phenol:chloroform:isoamyl and purified with ethanol precipitation and TE resuspension as above for ChIP DNA. qPCR was performed as above with the same primers to check *Airn* DNA enrichment.

CHART-Seq libraries were prepared as above for ChIP-Seq libraries. Single-end, 75-bp sequencing was performed using an Illumina NextSeq 500/550 High Output v2.5 kit on a NextSeq 500 System.

### RNA isolation, RT-qPCR, RNA-Seq

TSCs were passaged once off of irMEFs as above onto a single well of a 6-well plate. ESCs were grown on a single well of a 6-well plate. Both were grown to ≥75% confluency prior to RNA harvest using 1mL TRIzol, followed by the addition of 200μL chloroform, which were vortexed and subsequently spun at max speed for 5 minutes at 4°C for phase separation. The aqueous layer was collected and combined with 1 volume of 100% isopropanol and 5μL linear acrylamide. Precipitation was achieved at -80°C for 1 hour, followed by a max speed spin for 30 minutes at 4°C and one wash of the RNA pellet with ice-cold 80% ethanol. The pellet was then resuspended in 100μL H2O and quantified via Qubit (Invitrogen, cat #: Q32855).

For RT-qPCR assays in Figures 5A and S4E, 1μg of RNA was reverse transcribed using the High-Capacity cDNA Reverse Transcription Kit, and qPCR was performed using iTaq Universal SYBR Green Supermix and custom primers (Table S5).

RNA-Seq libraries were prepared from 1μg of total RNA using KAPA RNA HyperPrep Kit with Ribose Erase (Kapa Biosystems, cat #: KR1351) according to the manufacturer instructions. Single-end, 75-bp sequencing was performed using an Illumina NextSeq 500/550 High Output v2.5 kit on a NextSeq 500 System.

## DATA QUANTIFICATION AND STATISTICAL ANALYSIS

### Sequence alignment and processing

All mouse reference NCBI build 37/mm9 genome annotations were obtained from the UCSC genome browser (Lee et al. 2022). Variant sequence data was obtained from the Sanger Institute (http://www.sanger.ac.uk/resources/mouse/genomes/; (Keane et al. 2011). The CAST/EiJ (CAST) pseudogenome creation was performed as in (Calabrese et al. 2012; Calabrese et al. 2015). Hi-C reads were aligned using BWA as a part of the Juicer pipeline (Durand et al. 2016a). ChIP- and CHART-Seq reads were aligned using bowtie2 with default parameters (Langmead and Salzberg 2012). RNA-Seq reads were aligned using STAR with default parameters (Dobin et al. 2013).

For all Hi-C analyses in this study, read pairs that had a mapping quality greater than or equal to 10 were used (see Juicer section). For all ChIP-, CHART- and RNA-Seq analyses in this study, reads that had a mapping quality greater than or equal to 30 were extracted with samtools (Li et al. 2009), and allele-specific read retention (i.e., reads that overlap at least one B6 or CAST SNP) was performed as in (Calabrese et al. 2012; Calabrese et al. 2015) using a custom perl script (intersect_reads_snps16.pl: see github).

### Chromosome tiling density plots

For all chromosome-scale tiling density plots of allelic Hi-C viewpoint, CHART-Seq, allelic ChIP-Seq, and ESC Hi-C data in Figures 2-7, S2, S3, S5, reads were summed in 10kb bins across each chromosome. Read counts were then divided by the total number of reads in the dataset and divided by a million (i.e., RPM). For allelic data, binned counts were divided by the number of B6/CAST SNPs detected in the bin genomic coordinates (i.e., SNP-norm RPM). Finally, bins were averaged every 9 bins in 1bin increments. For allelic Hi-C viewpoint data, we excluded bins whose aggregate SNP-overlapping read count across merged Hi-C datasets fell within the bottom quintile relative to bins in the rest of the genome. The allelic data in this group of bins were too sparse to be interpreted with confidence. For the same reason, for allelic CHART- and ChIP-Seq data, only bins with greater or equal to 25 SNPs were plotted.

For ESC *Airn* CHART-Seq and H3K27me3 ChIP-Seq data in Figures 4, reads were summed in 40kb bins with 4kb tiling across each chromosome, as performed in (Schertzer et al. 2019a) for non-allelic data. All binned reads were normalized for alignability, where reads per bin were divided by the proportion of bases per bin that were uniquely alignable at 75bp resolution (i.e., the predominant read length used in this study) to avoid potential uncertainty that would be introduced by normalizing highly non-unique regions). Bins with alignability of less than 0.5 (i.e., less than 50% alignable at 75-bp resolution) were retained, and the read counts were divided by the total number of reads in the dataset and divided by a million (i.e., RPM).

All plots were generated using ggplot2 (Wickham 2016) in RStudio.

### Tiling density correlations

To derive Spearman’s ρ and p values for tiling density correlations in Figures 2 and 4, reads were processed as described in “Chromosome tiling density plots” and every 10^th^ bin along the genomic region of interest was correlated.

To determine significant changes in H3K27me3 and H2AK119ub density across genotypes in Figures 5 and S5, all binned SNP-norm RPM values over the genomic region of analysis were subjected to a Welch t-test.

### UCSC genome browser density tracks

Wiggle density files were created using a custom perl script (bigbowtie_to_wig3_mm9.pl; see github) and loaded into a UCSC Genome Browser session to generate the graphics in Figures 3 and 7. All density tracks were auto-scaled to data view and set to a maximum windowing function with 3-pixel smoothing.

### Hi-C analysis

#### Juicer processing, quality control, and allele-specific read retention

Hi-C analyses were carried out using a combination of Juicer (default parameters) and Hi-C Explorer (Durand et al. 2016a; Ramirez et al. 2018; Robinson et al. 2018). For exact commands used, see github.

For quality control, Hi-C statistics of each dataset were generated by Juicer (Table S1) and were referenced to the standard guidelines in (Rao et al. 2014). In addition, long-range DNA interactions (25kb-10Mb) were correlated between Hi-C replicate datasets at 10kb and 25kb resolutions using Hi-C Explorer’s hicCorrelate (Wolff et al. 2018) with the following parameters: --method=pearson --range 25000:10000000.

For allele-specific retention of Hi-C contacts, read pairs in which at least one read end overlaps a B6/CAST SNP were extracted from the Juicer output merged_nodups.txt file using a modified Juicer diploid script (juicer_diploid_v6.sh; see github).

#### 2D contact heatmaps

Allelic 2D contact heatmaps in Figures 1, 6, 7, S1, S6 were generated with Juicebox (Robinson et al. 2018). Contact matrices of observed counts were viewed in KR (Knight-Ruiz) balance mode at 5kb and 50kb resolutions. Subtraction heatmaps, where maternal contacts were subtracted from paternal contacts, were viewed under the same conditions.

#### Loop calling

DNA loops were detected using the Significant Interaction Peak (SIP) caller (Rowley et al. 2020) with the following parameters: -factor 4 -g 2.0 -t 2000 -fdr 0.05. The output finalLoops.txt file (i.e., a merged list of unique DNA loop anchors detected across 5k, 10kb, and 25kb resolutions) was used to determine loops over the 2D contact heatmaps in Juicebox for Figure 7.

“A” and “B” Compartmentalization

“A” and “B” chromosome compartments for each allele were delineated by eigenvector analysis using the Juicer *eigenvector* tool with the following parameters: -p KR 17 BP 50000. Extracted eigenvector values were then visualized and plotted with Juicebox in Figures 1 and 6.

#### Allelic viewpoints

Allelic viewpoints of locus-specific contact matrices were extracted at 10 or 25kb resolution for observed and observed-over-expected (O/E) counts using the Juicer *dump* tool with the following parameters: observed/oe NONE chr:start:end chr:start:end BP 10000/25000. If applicable, all viewpoint loci of interest were extended to 100kb lengths from their centers to improve contact matrix coverage. Extracted counts were then processed and plotted as described in the Chromosome-scale tiling density plots section.

#### Correlation with FISH

Allelic observed contacts from *Airn* viewpoint data in C/B wildtype TSCs were summed over the genomic coordinates for each of the 9 FISH probes across the *Airn* target domain analyzed in (Schertzer et al. 2019a). For each allele, the summed Hi-C counts at the probe locations were then correlated by Spearman’s test with the corresponding average distance to the *Airn* probe as measured by RNA/DNA co-FISH in C/B TSCs (Schertzer et al. 2019a). Scatter plots in Figure 2 were generated with ggplot2 in RStudio (Wickham 2016).

### ChIP-Seq analysis

#### Peak calling

ChIP-Seq peaks were called from non-allelic reads against an H3 ChIP-Seq dataset (from (Calabrese et al. 2012)) using the MACS2 peak calling algorithm (Zhang et al. 2008) with the following parameters: -g mm –broad –broad-cutoff 0.01.

#### Allelic enrichment at CpG islands and other features

A statistical permutation test was performed to determine how significantly enriched PRC components, CTCF, SMC1A/Cohesin, and epigenetic marks are at loci of interest relative to the rest of the genome (see Table S3). All datasets analyzed were generated from C/B TSCs. H3K27ac, H3K4me2, CTCF, and SMC1A data were generated in previous studies, as a part of (Calabrese et al. 2012; Schertzer et al. 2019a). If applicable, all genomic features of interest were standardized to 1.5kb lengths (i.e., the largest CGI of interest) relative to their center positions.

Using bedtools’ ‘shuffle’ (Quinlan and Hall 2010), we created a list of 80,000 1.5kb regions randomly selected from within ‘gene’ coordinates from gencode.vM1.annotation.gtf (Frankish et al. 2021) with 100kb extended start and end sites while excluding any regions that fell within 2.5kb of a region annotated by MACS as an H3K27me3 or PRC subunit peak. Shuffled coordinates were filtered to retain regions encompassing at least one B6/CAST SNP, leaving 67,262 shuffled regions. B6- and CAST-overlapping ChIP-Seq reads were then counted over the features of interest and shuffled coordinates using a custom script (ase_analyzer10_adj2.pl; see github), then divided by the number of B6/CAST SNPs detected in the genomic coordinates (SNP-norm counts). The features were then ranked by SNP-norm counts for each allele in each dataset (1 = highest allelic signal), and a percentile rank was used to determine an empirical p-value for allele-specific enrichment of the ChIP target at the loci of interest.

### RNA-Seq analysis

For non-allelic expression analysis in Table S2, featureCounts (Liao et al. 2014) was used to count reads over ‘gene’ entries in gencode.vM1.annotation.gtf (Frankish et al. 2021). Counts were then divided by total reads in the dataset and divided by a million (RPM).

For allelic expression analysis in Table S2 and Figures 5, S5, a custom perl script (ase_analyzer10.pl; see github) was used to count B6- and CAST SNP-overlapping reads over ‘gene’ entries in gencode.vM1.annotation.gtf (Frankish et al. 2021). Read counts were divided by the total number of reads in the dataset, divided by a million, and divided by the number of SNPs detected within the ‘gene’ coordinates (SNP-norm RPM). To determine the relative paternal expression of *Airn* in Figure S5E, paternal SNP-norm RPM values over *Airn* for each genotype were divided by was divided by the averaged NTG value from all four NTG clone data. To determine the relative paternal bias of *Airn* target gene expression for each genotype in Figure 5, paternal SNP-norm RPM values for the 27 *Airn* target genes (Table S2; (Schertzer et al. 2019a)) were divided by the sum of the paternal and maternal values. The paternal biases for each genotype were then divided by the averaged NTG value from all four NTG clone data, and a Welch t-test was used to determine a statistical p-value of the relative change in expression relative to NTG. Boxplots of these data were generated using GraphPad Prism v9.

